# TRPA1 analgesia is mediated by kappa opioid receptors

**DOI:** 10.1101/2022.09.01.506151

**Authors:** Evangelia Semizoglou, Clive Gentry, Nisha Vastani, Cheryl L. Stucky, David A. Andersson, Stuart Bevan

## Abstract

TRPA1 expressed in peripheral sensory neurons is important for nociception. Pharmacological inhibition or genetic ablation of TRPA1 profoundly reduces normal behavioural sensitivity to noxious cold and mechanical stimulation, as well as sensory neuron responses to mechanical stimulation. TRPA1 inhibition also reverses cold and mechanical hypersensitivities in chronic pain models *in vivo*. Here we demonstrate that these striking effects of TRPA1 inactivation result from an increased constitutive activity of kappa opioid receptors (KOR) co-expressed with TRPA1 in sensory neurons. Inhibition of KOR in *Trpa1^-/-^* mice restores nociception and neuronal activity to the levels observed in wild-type mice and reverses the analgesic effects of TRPA1 antagonism in naïve mice and in neuropathic and inflammatory pain conditions. TRPA1 regulation of KOR activity in sensory neurons provides a novel mechanism to produce peripherally mediated analgesia. Our findings suggest that TRP channel regulation of constitutive GPCR activity, may be a process of general physiological importance.

## INTRODUCTION

The ion channel TRPA1 is highly expressed in small to medium sized, unmyelinated or lightly myelinated dorsal root ganglion (DRG) neurons, many of which co-express TRPV1 and CGRP (Story et al., 2003, Li et al., 2016, Brierley et al., 2011, Jordt et al., 2004). Here, it acts as a sensor of many noxious chemicals and is activated by a range of cysteine-modifying oxidants and electrophiles, including pungent exogenous agents such as allyl isothiocyanate (AITC) and tear gases, as well as endogenous electrophiles such as reactive oxygen species, methylglyoxal and lipid peroxidation products (see Talavera et al., 2020, Viana, 2016).

TRPA1 has also been implicated in the regulation of thermal and mechanical sensitivities in mammals although the underlying mechanisms are unknown. TRPA1 was originally reported as a sensor of noxious cold (Story et al., 2003), but its role as a primary cellular cold transducer is contentious (for a recent review see Talavera et al., 2020). Behavioural studies of mice lacking functional TRPA1 (*Trpa1^-/-^* mice) have similarly reported conflicting data on the role of TRPA1 in cold sensation *in vivo* (Bautista et al., 2006, del Camino et al., 2010, Gentry et al., 2010, Knowlton et al., 2010, Kwan et al., 2006, Zappia et al., 2017). Although there is no strong evidence that TRPA1 is a bona fide mechanotransduction channel in cells, it has also been implicated in the regulation of mechanosensation. For example, *Trpa1^-/-^* mice display reduced sensitivity to noxious mechanical stimulation in behavioural tests (Andersson et al., 2009, Kwan et al., 2006, Zappia et al., 2017). Furthermore, electrophysiological studies of intact skin-nerve preparations identified reduced mechanically evoked impulse activity in preparations from *Trpa1^-/-^* mice, as well as in wild-type mice treated with a TRPA1 antagonist (Kerstein et al., 2009, Kwan et al., 2009, Vilceanu and Stucky, 2010).

In the current study we have investigated the mechanisms by which genetic ablation and pharmacological inhibition of TRPA1 influence responses to mechanical and cold stimuli. Importantly, our data reveal that the sensory deficits observed in *Trpa1^-/-^* mice are due to increased constitutive kappa opioid receptor (KOR) activity in sensory neurons. Our studies further show that TRPA1 antagonists produce profound antinociception in naïve animals by generating constitutive KOR activity acutely, which reduces sensory neuron firing. Intriguingly these results demonstrate that the activity of an ion channel, TRPA1, regulates the activity of a G protein coupled receptor. This novel KOR-dependent mechanism also operates in mouse models of neuropathic and inflammatory pain and explains the general analgesic effects of TRPA1 deletion and TRPA1 antagonism.

## RESULTS

### TRPA1 regulates cold and mechanical sensitivity

We examined behavioural responses of *Trpa1^-/-^* and wild-type mice to thermal stimulation over the range of temperatures from 2°C to 55°C using a modified cold/hot plate assay, to determine the behavioural importance of TRPA1 for thermal nociception. In this assay, the paw is in constant contact with the temperature-controlled plate and the end-point is clear and robust (Gentry et al., 2010). Previous studies on cold sensitivity have used different strains of *Trpa1^-/-^* mice, which may have contributed to divergent results, and we therefore compared responses in mice generated in the Kwan/Corey (*Trpa1^-/-Kykw^*) (Kwan et al., 2006) and Julius (*Trpa1^-/-Jul^*) (Bautista et al., 2006) laboratories. The behavioural profiles of the two *Trpa1^-/-^* strains were indistinguishable (Fig. 1a). We observed no sex differences in these behavioural tests and data from male and female mice have therefore been amalgamated. *Trpa1^-/-^* mice exhibited a pronounced decrease in cold sensitivity compared to wild-type *Trpa1^+/+^* mice at temperatures between 5 and 20°C. The sensitivity to the coldest temperature tested (2°C) was, however, identical in the two genotypes (Fig 1a). Examination of the responses evoked by a 10°C stimulus in *Trpa1^+/+^, Trpa1^-/-^* and heterozygous (*Trpa1^+/-^*) mice further revealed a gene dosage effect of TRPA1 on withdrawal latency (*Trpa1^-/-^* > *Trpa1^+/-^ > Trpa1^+/+^*, Fig 1b). In contrast to the effects on cold nociception, the paw withdrawal latencies at higher temperatures (25-55°C) were identical in *Trpa1^-/-^*, Trpa1^+/-^ and wild-type (*Trpa1^+/+^*)^-^ mice (Fig 1a and 1c, 53°C stimulus).

**Figure 1.**
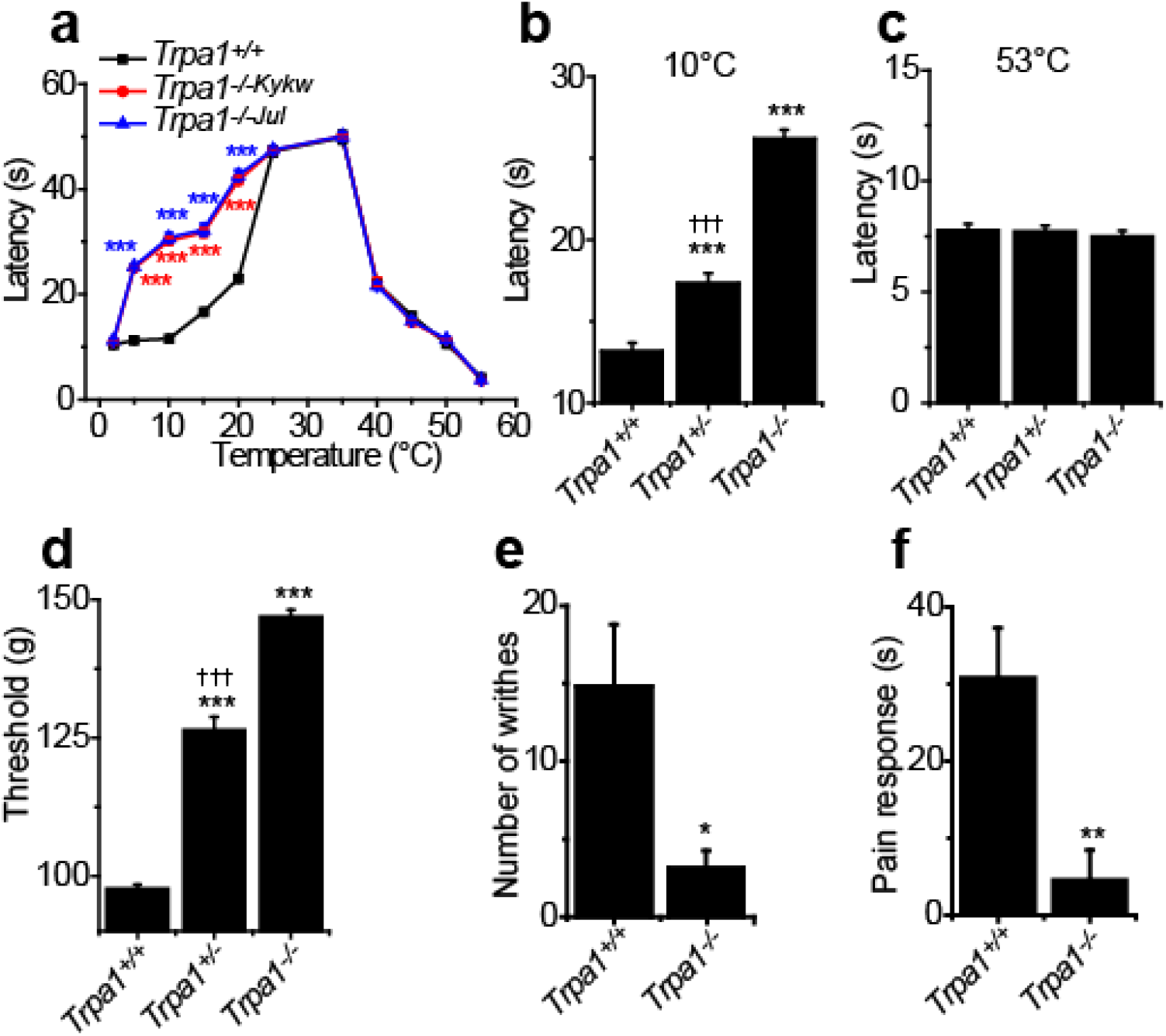
TRPA1 controls noxious cold and mechanical sensitivity. (a) Two independent strains of *Trpa1^-/-^* mice (*Trpa1^-/-Kykw^* and *Trpa1^-/-Jul^*) display an identical and significantly increased paw withdrawal latency compared to wild-type mice in the temperature range 5-20°C (using the cold/hot plate test). (b) Responsiveness to a 10°C cold plate is regulated by TRPA1 expression, whereas heat nociception (c) is unaffected by *Trpa1* status. (d) Paw withdrawal threshold in the paw pressure test is regulated by TRPA1. Abdominal constrictions induced by intraperitoneal phenyl-*p*-benzoquinone (e) and paw licking, biting and flinching induced by intraplantar AITC (f) are significantly reduced in *Trpa1^-/-^* compared to *Trpa1^+/+^* mice. *P<0.05, **P<0.01 and ***P<0.001, compared to *Trpa1^+/+^* mice; ††† P<0.001, compared to *Trpa1^-/-^* mice; repeated measures two-way ANOVA with Sidak’s post-hoc test (a), ANOVA followed by Tukey’s post-hoc test (b-d), or unpaired t-test (e, f). Each data point is the mean ± SEM of at least 7 mice.

We next examined mechanical nociception in *Trpa1^-/-^, Trpa1^+/-^ and Trpa1^+/+^* mice, using the Randall-Selitto paw pressure test. As previously reported (Kwan et al., 2006, Andersson et al., 2009), TRPA1 exerted a strong influence over mechanical nociception and, once again, the loss of sensitivity followed a gene-dosage relationship (*Trpa1^-/-^* > *Trpa1^+/-^ > Trpa1^+/+^*, Fig 1d). As expected from the role of TRPA1 as a sensory receptor for reactive chemical irritants, *Trpa1^-/-^* mice showed a loss of chemosensitivity in behavioural tests measuring responses to either intraperitoneal administration of the electrophilic TRPA1 agonist phenyl-p-benzoquinone (Fig 1e) or intraplantar injection of AITC (Fig 1f).

### Opioid receptors mediate TRPA1 antinociception

Given the marked effects of TRPA1 inactivation on cold and mechanical sensitivities, which are not directly transduced by TRPA1, we asked whether the reduction in sensitivities could be mediated indirectly by endogenous analgesic mechanisms. Opioid receptor activity masks long-lasting hypersensitivities that continue after inflammation or tissue injury, and administration of opioid antagonists can unmask this latent sensitization to reveal the underlying hypersensitivities (Campillo et al., 2011, Corder et al., 2013, Custodio-Patsey et al., 2020, Severino et al., 2018, Walwyn et al., 2016). We therefore investigated the effects of the broad-spectrum opioid receptor antagonist naloxone on the behavioural responses of *Trpa1^-/-^* mice. Systemic administration of naloxone (i.p. injection) dose-dependently normalised the mechanical responsiveness in both strains of *Trpa1^-/-^* mice, reducing the paw pressure thresholds to the level of wild-type mice (Fig 2a and Supplementary Fig 1a, *Trpa1^-/-Kykw^* and Supplementary Fig 1c, *Trpa1^-/-Jul^*) but was without effect in wildtype mice. Similarly, the reduced sensitivity of *Trpa1^-/-^* mice in a 10°C cold plate test was normalised by naloxone (Fig 2b and Supplementary Fig 1b, *Trpa1^-/-Kykw^* and Supplementary Fig. 1d, *Trpa1^-/-Jul^*). Next, we explored the effects of naloxone in *Trpa1^-/-^* mice over a range of cold temperatures (2-25°C, Fig 2c). Naloxone reduced the paw withdrawal latency to values observed in wild-type mice at temperatures where there was a difference between *Trpa1^-/-^* and wild-type mice (5-20°C) but had no effect on response latencies at temperatures that were unaffected by the TRPA1 status (2 and 25°C). In contrast to its effects in *Trpa1^-/-^* mice, naloxone had no discernible effect on the cold sensitivity of *Trpa1^+/+^* mice at any temperature tested (Fig 2c).

**Figure 2.**
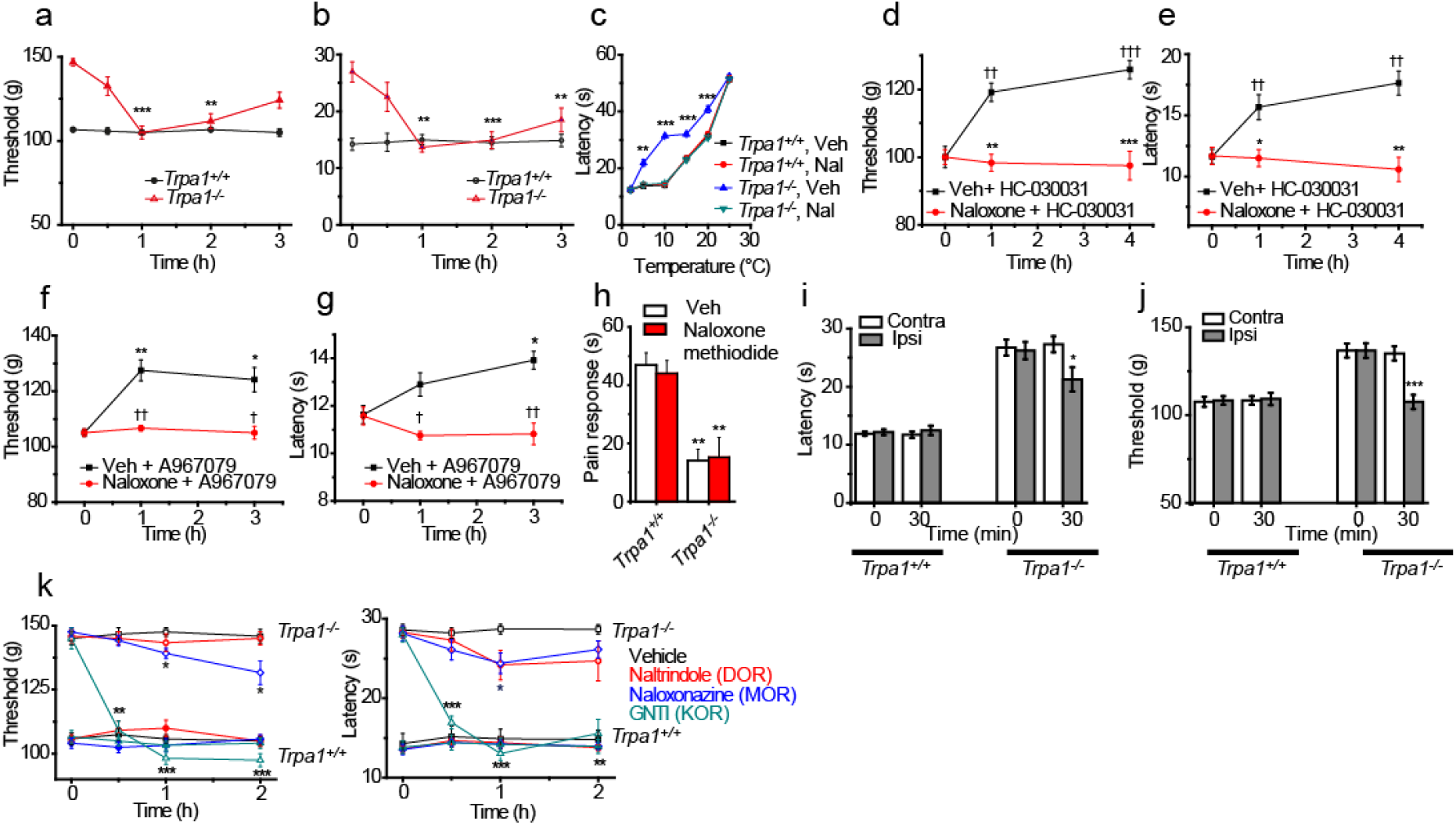
Opioid receptors mediate the antinociceptive effect of TRPA1 inactivation. Naloxone (2.5mg/kg, i.p.) reduces paw-pressure withdrawal threshold (a) and cold-plate latency (b, 10°C) in *Trpa1^-/-^* but not *Trpa1^+/+^* mice. (c) Effect of naloxone on the thermal sensitivity of *Trpa1^-/-^* and *Trpa1^+/+^* mice in the temperature range 2-25°C. TRPA1 antagonists HC-030031 (d, e) and A967079 (f, g) increased the paw-pressure withdrawal threshold and cold-plate latency in wild-type mice. Pre-treatment with naloxone abolished the antinociceptive effect of HC-030031 (d, e) and A967079 (f, g). Intraplantar AITC (100μmoles in 25μl) evoked similar pain-related behaviour in wild-type mice treated with vehicle or naloxone methiodide (2.5mg/kg, i.p.). (h) The nocifensive response evoked by AITC in *Trpa1^-/-^* mice was markedly reduced and unaffected by naloxone methiodide. Intraplantar injection of naloxone methiodide reduced the ipsilateral cold-plate paw withdrawal latency (i) and paw pressure withdrawal threshold (j) in *Trpa1^-/-^*, but was without effect in wild-type mice and in the contralateral paw of *Trpa1^-/-^* mice. (k) Paw pressure withdrawal latency and (l) cold plate latency in *Trpa1^-/-^* mice (empty symbols) were significantly reduced compared to vehicle (0.2ml saline, s.c., black squares) by administration of the KOR antagonist GNTI (0.3 mg/kg, s.c., cyan triangles). The MOR antagonist naloxonazine (35mg/kg, s.c., blue diamonds) and DOR antagonist naltrindole (5mg/kg, s.c., red circles), had little or no effect on *Trpa1^-/-^* paw pressure withdrawal threshold and cold plate latency. The responsiveness of wild-type mice was unaffected by all treatments (filled symbols). *P<0.05, **P<0.01 and ***P<0.001, effect of naloxone compared to vehicle, treatment or naïve pre-treatment reading, repeated measures two-way ANOVA followed by Dunnett’s or Sidak’s post-hoc tests, or unpaired t-test. †† P<0.01, ††† P<0.001, effect of TRPA1 antagonist treatment. Each data point is the mean ± SEM of at least 6 mice.

To assess whether the opioid-dependent antinociception observed in *Trpa1-/-* mice was a consequence of the global, lifelong TRPA1 inactivation, or a more dynamic, ongoing process, we determined whether the effects of two structurally different TRPA1 antagonists were naloxone sensitive. Administration of a single dose of either HC-030013 (Fig. 2d, e) or A-967079 (Fig. 2f, g) to wild-type mice reduced their sensitivities to paw pressure and cold plate stimuli, as shown by elevated pressure thresholds and prolonged cold withdrawal latencies, thereby recapitulating the nociceptive phenotype observed in *Trpa1^-/-^* mice. These effects of the TRPA1 antagonists were abolished by prior administration of naloxone, indicating that TRPA1 antagonism reduced sensitivities to noxious cold and mechanical stimulation by engaging an opioid receptor mediated pathway acutely.

### A peripheral site for opioid receptor action

Opioids can produce analgesia by stimulating receptors expressed in the central nervous system (CNS) or located on peripheral sensory neurons. We therefore investigated the site of action of naloxone in *Trpa1^-/-^* mice using the peripherally restricted analogue naloxone methiodide. First, we tested the effects of systemic naloxone methiodide on the behavioural response to direct activation of TRPA1 elicited by intraplantar injections of the TRPA1 agonist AITC. AITC evoked a pronounced TRPA1-mediated pain response (flinching, licking) in wild-type mice that was unaffected by naloxone methiodide, demonstrating that the opioid antagonist did not interfere with TRPA1-dependent nociception. As expected for a TRPA1 agonist, the reduced sensitivity of *Trpa1^-/-^* mice to AITC was not reversed by naloxone methiodide Fig 2h).

Next, we investigated the effect of intraplantar injection of naloxone methiodide on cold (Fig 2i) and mechanical (Fig 2j) sensitivities in *Trpa1^+/+^* and *Trpa1^-/-^* mice. Naloxone methiodide significantly reduced the *Trpa1^-/-^* ipsilateral paw withdrawal latency in the cold plate test and the withdrawal threshold in the paw-pressure test to the levels observed in *Trpa1^+/+^* mice. However, as observed with systemically active naloxone, intraplantar naloxone methiodide had no effect in *Trpa1^+/+^* mice. Importantly, this treatment had no effect on the cold latency or mechanical threshold in the contralateral paw, demonstrating that the actions of naloxone methiodide were limited to the treated paw. These results indicate that the opioid pathway-mediated effects in *Trpa1^-/-^* mice operate locally in the injected paw, rather than in the CNS.

### Kappa opioid receptors are responsible for the deficits in Trpa1^-/-^ mice

To identify the opioid receptor sub-type responsible for the reduced sensitivities in *Trpa1^-/-^* mice we administered selective antagonists for mu (MOR), delta (DOR) and kappa (KOR) opioid receptors. The KOR antagonist 5’-GNTI completely reversed the reduced mechanical (Fig 2k) and cold (Fig 2l) sensitivities in *Trpa1^-/-^* mice, restoring the sensitivities to values observed in wild-type mice. In contrast, selective MOR (naloxonazine) and DOR (naltrindole) antagonists had little to no effect in *Trpa1^-/-^* mice. The behavioural nociceptive responses of *Trpa1^+/+^* mice were also unaffected by these opioid receptor antagonists (Fig 2k,l).

### A TRPA1 - opioid receptor interaction operates in chronic pain models

TRPA1 has attracted attention as an analgesic target since selective antagonists inhibit painful hypersensitivities in multiple experimental pain models. Therefore, we examined the cold and mechanical sensitivity of *Trpa1^-/-^* and *Trpa1^+/+^* mice in a partial nerve ligation (PNL) model of traumatic neuropathic pain. As expected, the pre-surgery baseline cold withdrawal latencies and mechanical thresholds were elevated in *Trpa1^-/-^* mice. After surgery, the cold latency (Fig 3a) and mechanical threshold (Fig 3b) were reduced in both genotypes by day 14, although the sensitivities in *Trpa1^-/-^* mice remained higher than in naive *Trpa1^+/+^* mice. Strikingly, naloxone increased the cold and mechanical sensitivities of the operated paw in *Trpa1^-/-^* mice to the same hypersensitive levels observed in neuropathic *Trpa1^+/+^* mice. Naloxone had no effect on the hypersensitivities in *Trpa1^+/+^* mice. As already noted in naïve unoperated mice, the cold and mechanical sensitivities of the contralateral, uninjured paws of *Trpa1^-/-^* mice increased to the levels recorded in *Trpa1^+/+^* mice.

**Figure 3.**
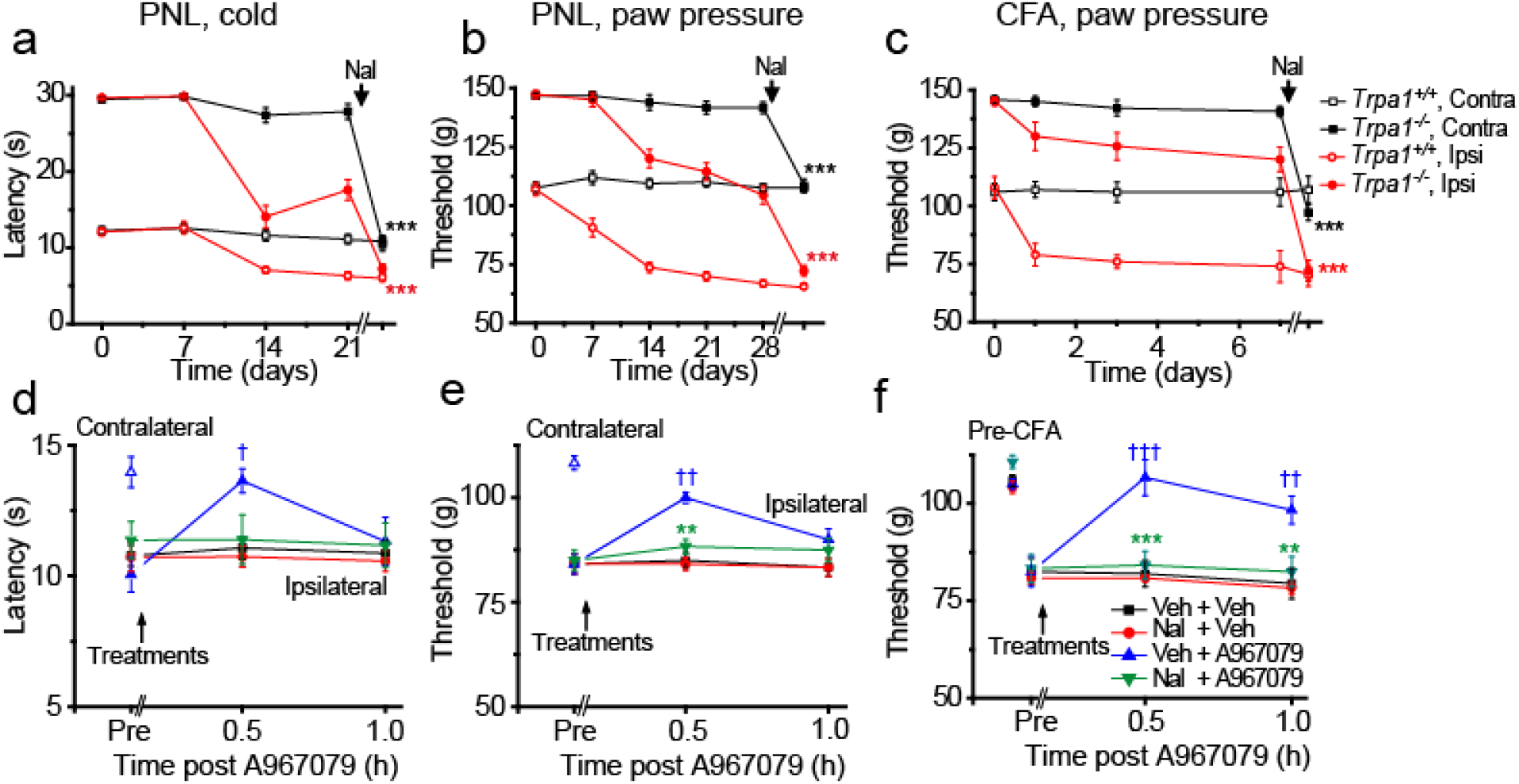
Opioid receptors mediate the analgesic effect of TRPA1 inactivation in chronic pain models. Partial ligation of the sciatic nerve (PNL) increases the ipsilateral sensitivity of *Trpa1^+/+^* and *Trpa1^-/-^* mice in the cold-plate (a) and paw pressure tests (b). Following administration of naloxone (2.5mg/kg, i.p.), both the ipsi- and contralateral sensitivity of *Trpa1^-/-^* mice were increased to the wild-type level (1h post naloxone). Note that the effect of naloxone (c) Intraplantar injections of complete Freund’s adjuvant (CFA), reduced the ipsilateral paw withdrawal threshold in the paw-pressure test in *Trpa1^-/-^* and *Trpa1^+/+^* mice. Naloxone increased the *Trpa1^-/-^* sensitivity to the same level as the *Trpa1^+/+^* mice. ***P<0.001 vs pre naloxone administration, t-test, 8-9 mice in each group. Note that naloxone was without effect on the neuropathic or inflammatory hypersensitivities in *Trpa1^+/+^* mice. Naloxone inhibits the A967079-mediated reversal of established neuropathic cold (d) and paw pressure (e) hypersensitivities in PNL wild-type mice. (f) Naloxone also inhibited the anti-hyperalgesic effect of A987079 on paw pressure sensitivity after establishment of CFA-induced inflammation. The effects of A967079 on the contralateral untreated paws in PNL and CFA mice are consistent with data in Fig. 2f,g. †P<0.05 and †††P<0.001, effect of A967079 compared to pre-treatment. **P<0.01, ***P<0.001 effect of naloxone compared with A967079 alone: repeated measure two-way ANOVA with Dunnett’s or Sidak’s post hoc tests. 6 mice per group. Each data point is the mean ± SEM.

Similar effects of naloxone were obtained in an inflammatory (Complete Freund’s Adjuvant, CFA) pain model (Fig 3c). Hypersensitivity to mechanical (paw pressure) stimulation developed in both *Trpa1^+/+^* and *Trpa1^-/-^* mice although the threshold for paw withdrawal in *Trpa1^-/-^* mice remained higher than in *Trpa1^+/+^* mice. Once again, naloxone increased the mechanical sensitivities of *Trpa1^-/-^* inflamed and non-inflamed paws to the corresponding levels observed in CFA treated *Trpa1^+/+^* mice (Fig 3c). Naloxone therefore unmasked a peripheral analgesic opioid system mechanism that still operates in *Trpa1^-/-^* mice in chronic neuropathic and inflammatory pain conditions.

### Naloxone reverses the effects of a TRPA1 antagonist in chronic pain models

Next, we investigated the effects of naloxone on the actions of the TRPA1 inhibitor A967079 in two models of chronic pain. Inflammatory and neuropathic mechanical and cold hypersensitivities were established by intraplantar injection of CFA or PNL surgery, respectively. Pronounced mechanical hypersensitivity was evident at 3 days in the CFA model when systemic administration of A967079 (30mg/kg p.o.) effectively reversed the hypersensitivity. However, the effect of A967079 was completely inhibited by prior treatment with naloxone, which alone had no effect on the behavioural response (Fig 3d). Similarly, naloxone inhibited the ability of A967079 to reverse established mechanical- and cold-hypersensitivities in the PNL model of neuropathic pain (Fig 3e,f).

### Sensory neuron TRPA1 determines the Trpa1^-/-^ phenotype

TRPA1 expression has been reported in tissues other than sensory neurons including astrocytes and Schwann cells (De Logu et al., 2017, Shigetomi et al., 2011). We therefore examined whether selective deletion of TRPA1 in sensory neurons was sufficient to recapitulate the naloxone-reversible loss of sensitivity seen in the global TRPA1 knockout mice. First, mechanical (paw pressure) and cold sensitivities were compared in Avil-Cre^+/-^ */Trpa1^fl/fl^* mice, where TRPA1 is selectively deleted from the sensory neurons (Zappia et al., 2017), and in global *Trpa1^-/-^* and control Avil-Cre^+/-^/*Trpa1^+/+^* mice. The mechanical thresholds of *Avil-Cre^+/-^/Trpa1^fl/fl^* mice were raised to the cut off value of 150g, similar to the profile observed in *Trpa1^-/-^* mice (Fig 4a). *Avil-Cre^+/-^/Trpa1^fl/fl^* mice also displayed a loss of cold sensitivity similar to that in *Trpa1^-/-^* mice (Fig 4b). The mechanical and cold thresholds in control Avil-*Cre^+/-^/Trpa1^+/+^* mice were similar to the values previously reported in wild-type mice (Fig 4a,b).

**Figure 4.**
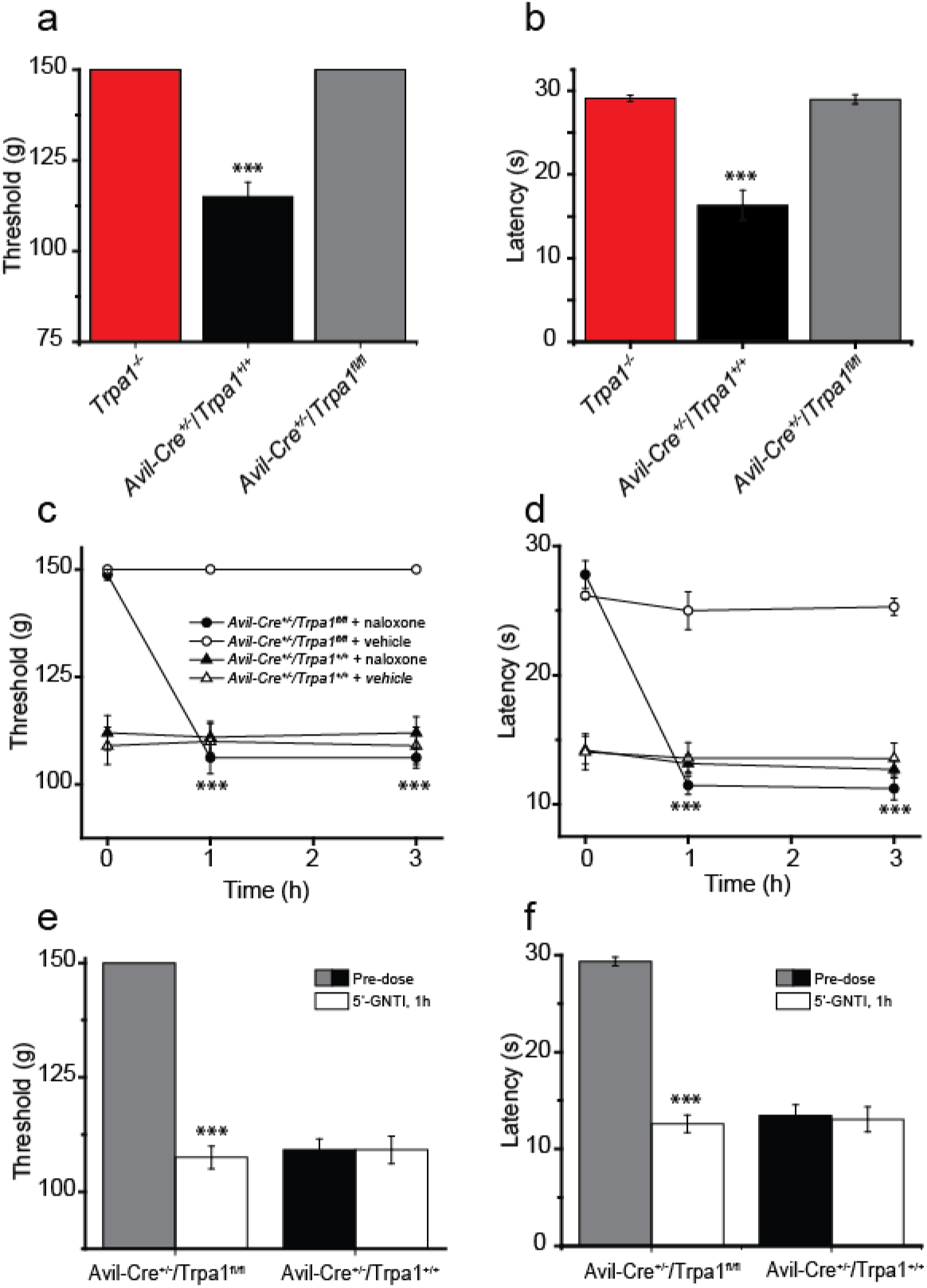
Sensory neuron TRPA1 determines the *Trpa1^-/-^* phenotype. (a) Paw pressure thresholds and (b) cold withdrawal latencies are increased in *Avil-Cre^+/-^-Trpa1^fl/fl^* and *Trpa1^-/-^* compared to *Avil-Cre^+/-^-Trpa1^+/+^* mice. Naloxone (2.5mg/kg, i.p.) increased mechanical (c) and cold (d) sensitivities in *Avil-Cre^+/-^-Trpa1^fl/fl^* to the levels observed in *Avil-Cre^+/-^-Trpa1^+/+^* mice. The KOR specific antagonist 5’-GNTI increases (e) mechanical and (f) cold sensitivities in *Avil-Cre^+/-^-Trpa1^fl/fl^* mice to the levels recorded in *Avil-Cre^+/-^-Trpa1^+/+^* mice, I hour after administration, 0.3 mg/kg, s.c. *** P<0.001, 6 mice per group, repeated measures two-way ANOVA followed by Dunnett’s or Sidak’s post-hoc tests.

Next, we determined the effects of naloxone and the KOR-specific antagonist 5’-GNTI on mechanical (Fig 4c, e) and cold (Fig 4d, f) responses. Neither compound had any effect on mechanical or cold responses in control Avil^+/*Cre*^/*Trpa*^+/+^ mice. In contrast, both compounds normalized the response thresholds in Avil^+/*Cre*^/*Trpa1^fl/fl^* mice, thereby demonstrating that the KOR-mediated normalization of cold and mechanical sensitivities is due to a loss of TRPA1 in the sensory neurons.

### TRPA1 and KOR are co-expressed in a sub-population of mouse DRG neurons

We initially determined the expression levels of *Oprk1* and *Trpa1* in *Trpa1^+/+^* and *Trpa1^-/-^* mouse DRGs using RT-PCR. *Trpa1* cDNA was targeted at two different regions, the channel S5 and S6 transmembrane domains (exons 21-25), which have been deleted in the *Trpa1^-/-^* mice (Kwan et al., 2006) and the ankyrin repeat region (exons 7-8), which is present in both *Trpa1^+/+^* and *Trpa1^-/-^* genomes. As expected, the signal obtained with the probe for the S5-S6 pore region in DRG neurons from *Trpa1^-/-^* mice was <0.2% of the level seen in wild-type mice, with a reduction (~10-fold) when a probe for the non-deleted ankyrin repeat sequence was used. No significant difference in *Oprk1* expression was detected between *Trpa1^+/+^* and *Trpa1^-/-^* DRGs (Fig. 5a).

**Figure 5.**
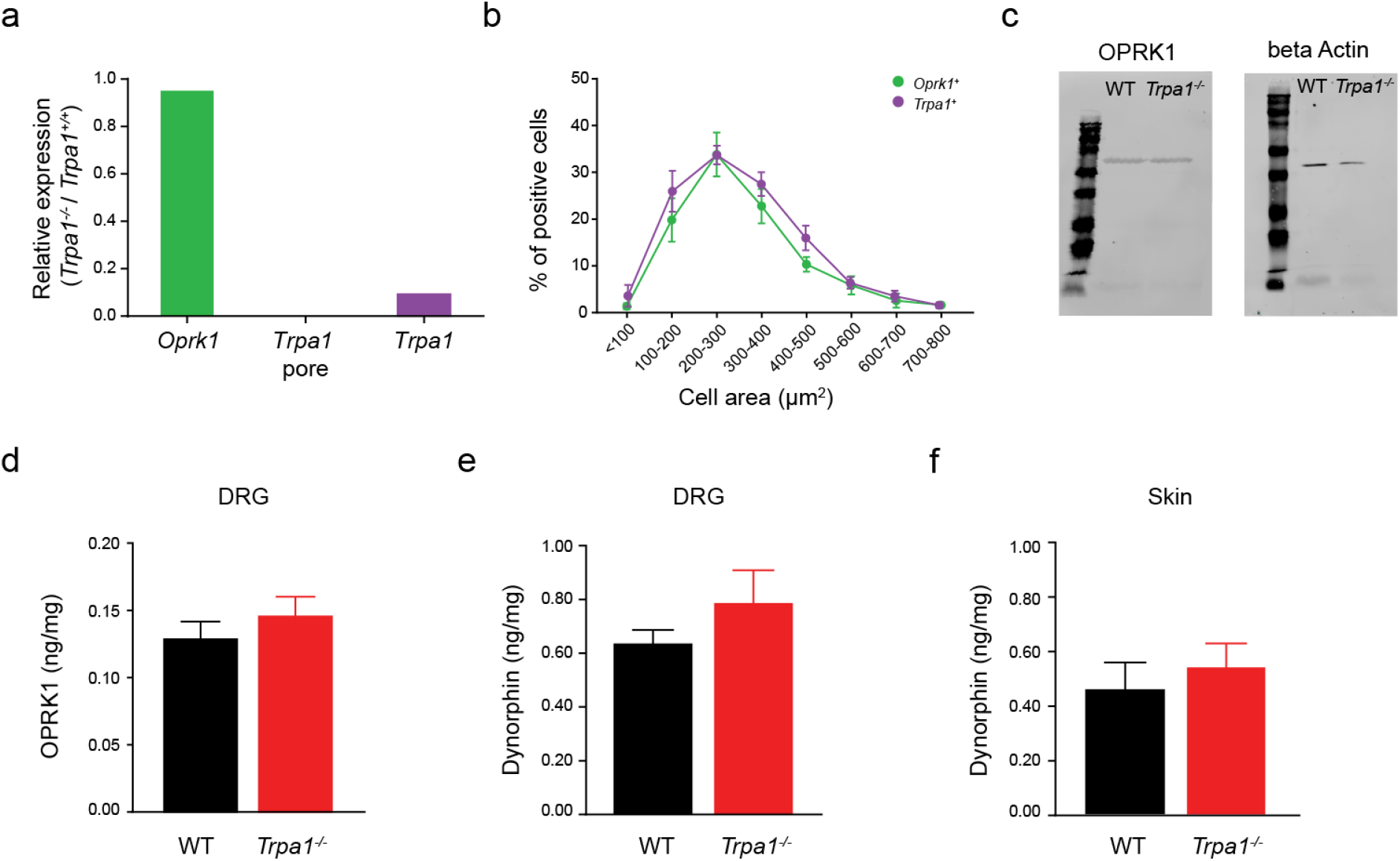
Expression of *Oprk1* mRNA, KOR protein and dynorphin in *Trpa1^-/-^* mice. (a) Relative expression levels of mRNA transcripts encoding *Oprk1*, the deleted S5-S6 pore region of *Trpa1* (Trpa1 pore) and the non-deleted ankyrin repeat region (*Trpa1*-ankyrin) in *Trpa1^-/-^* compared to *Trpa1^+/+^* mice. (b) Size distribution of DRG neurons expressing *Oprk1* (green) and *Trpa1* (purple). (c) Western blot of KOR protein in *Trpa1^+/+^* and *Trpa1^-/-^* mice (left), beta-actin expression used as protein loading control (right). ELISA measurements showing no significant difference in expression levels in *Trpa1^+/+^* and *Trpa1^-/-^* mice of (d) KOR in DRG (P=0.54) and dynorphin in (e) DRG (P=0.34) and (f) skin (P=0.59, n=6 for all samples, unpaired t-test).

To determine the expression pattern of Oprk1 and TRPA1 in DRG neurons, we examined the expression of *Oprk1* by *in situ* hybridization (ISH) using a RNAscope multiplex fluorescent assay (Wang et al., 2012) and compared the expression with the antibody staining for two neuronal markers, neurofilament 200 and the neuropeptide calcitonin gene related peptide (CGRP). *Oprk1* was expressed in both neurofilament-positive neurons and CGRP expressing neurons (Fig. 6a). *Oprk1* and *Trpa1* transcripts were expressed in small- to medium-diameter neurons (area 100-500μm^2^, diameter 10-25μm, Fig 5b). The expression pattern of *Oprk1* was unaffected by the Trpa1 genotype, and we detected a similar percentage of *Oprk1* expressing neurons in DRGs from *Trpa1^+/+^* (10.0%) and *Trpa1^-/-^* (11.4%) mice. Importantly, we noted an extensive co-expression of *Oprk1* and *Trpa1* with the *Trpa1* transcript expressed in similar percentages of *Oprk1*-expressing DRG neurons (*Trpa1^+/+^*, 42% and *Trpa1^-/-^*, 35%). This co-localization of *Trpa1* and *Oprk1* is consistent with a sensory neuron locus for the behavioural effects of KOR inhibitors in *Trpa1^-/-^* mice.

**Figure 6.**
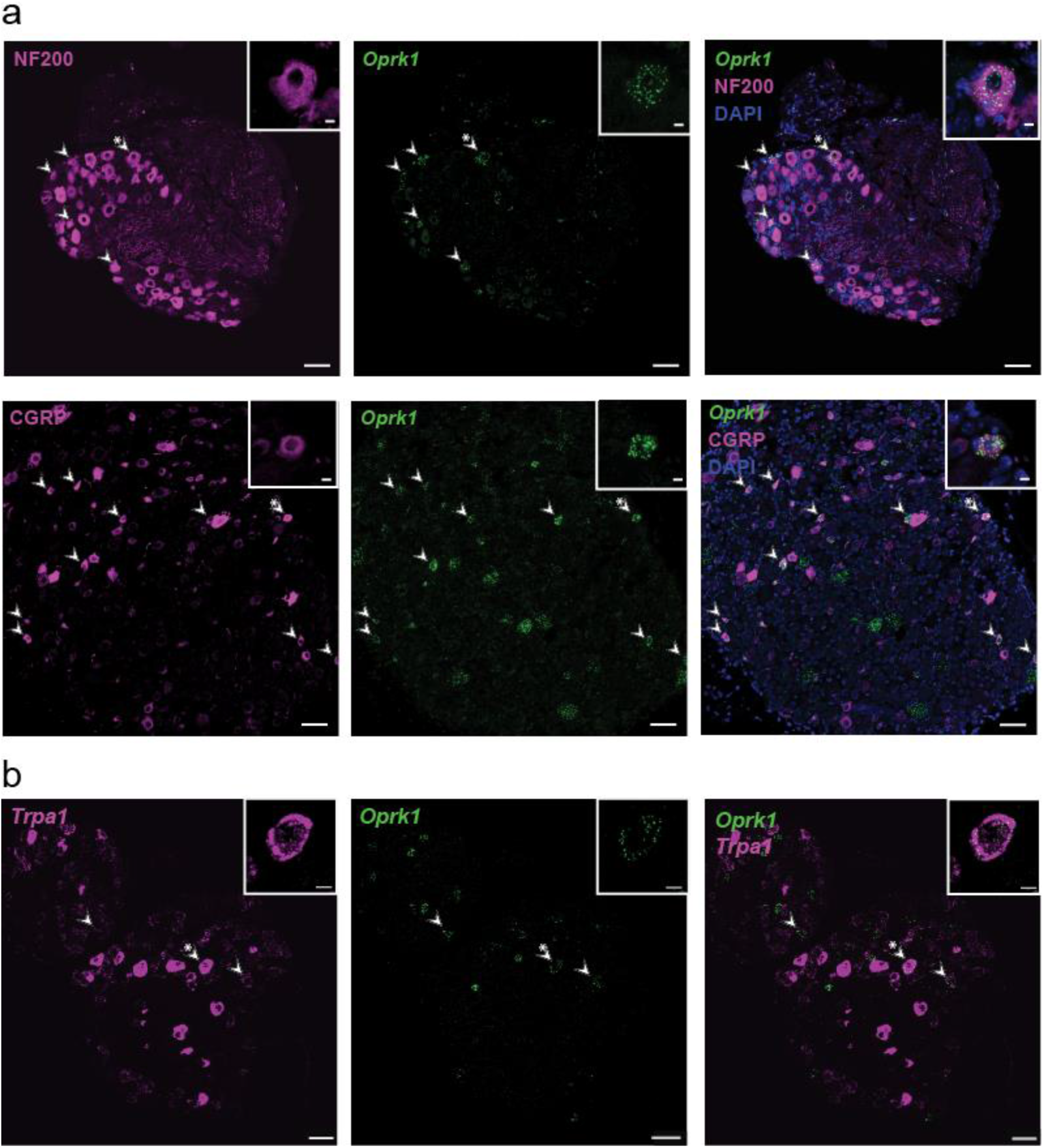
Co-expression of TPRA1 and KOR in a sub-population of DRG neurons. Combined *in situ* hybridization-immunostaining showing (a) co-expression of Oprk1 mRNA (green) and (top) neurofilament 200 (purple) and (bottom) CGRP (purple). (b) In situ hybridization of *Oprk1* mRNA (green) and *Trpa1* mRNA (purple). Arrowheads mark Oprk1-positive neurons co-expressing the other marker. Neurons shown in insets marked with asterisk. Bars 50 μm (x10) and 5μm (x40 insets).

As mRNA transcript expression may not reflect the degree of protein expression accurately, we also examined the expression of KOR protein in *Trpa1^-/-^* and *Trpa1^+/+^* mouse DRGs by western blotting and ELISA. No difference in DRG KOR protein expression was noted between the two genotypes using either technique (Fig. 5 c,d). We also assessed the possibility that increased levels of the endogenous agonist, dynorphin, were responsible for the increased KOR activity in *Trpa1^-/-^* mice. However, ELISA measurements of dynorphin levels in DRG neurons and skin, revealed no significant differences in dynorphin concentrations between *Trpa1^-/-^* and *Trpa1^+/+^* mice (Fig. 5 e,f). This is consistent with our findings that analgesia evoked by TRPA1 antagonists can be reversed by naloxone as these acute treatments would not be expected to change dynorphin levels.

### TRPA1 and KOR are co-expressed in human DRG neurons

To determine the translational relevance of our findings, we examined whether human DRG neurons co-expressed *TRPA1* and *OPRK1* using RNAscope ISH. *TRPA1* mRNA was detected in 29% of neurons, which is somewhat higher than the percentage (16%) reported in a recent ISH study (Tavares-Ferreira et al., 2022). Notably, TRPA1 was clearly co-localized with *OPRK1* in some neurons (Fig. 7). However, the relatively low level of *OPRK1* mRNA expression in individual neurons precluded quantitative studies of the degree of co-expression. Nevertheless, this co-localization raises the possibility that TRPA1 regulation of KOR activity also occurs in humans.

**Figure 7.**
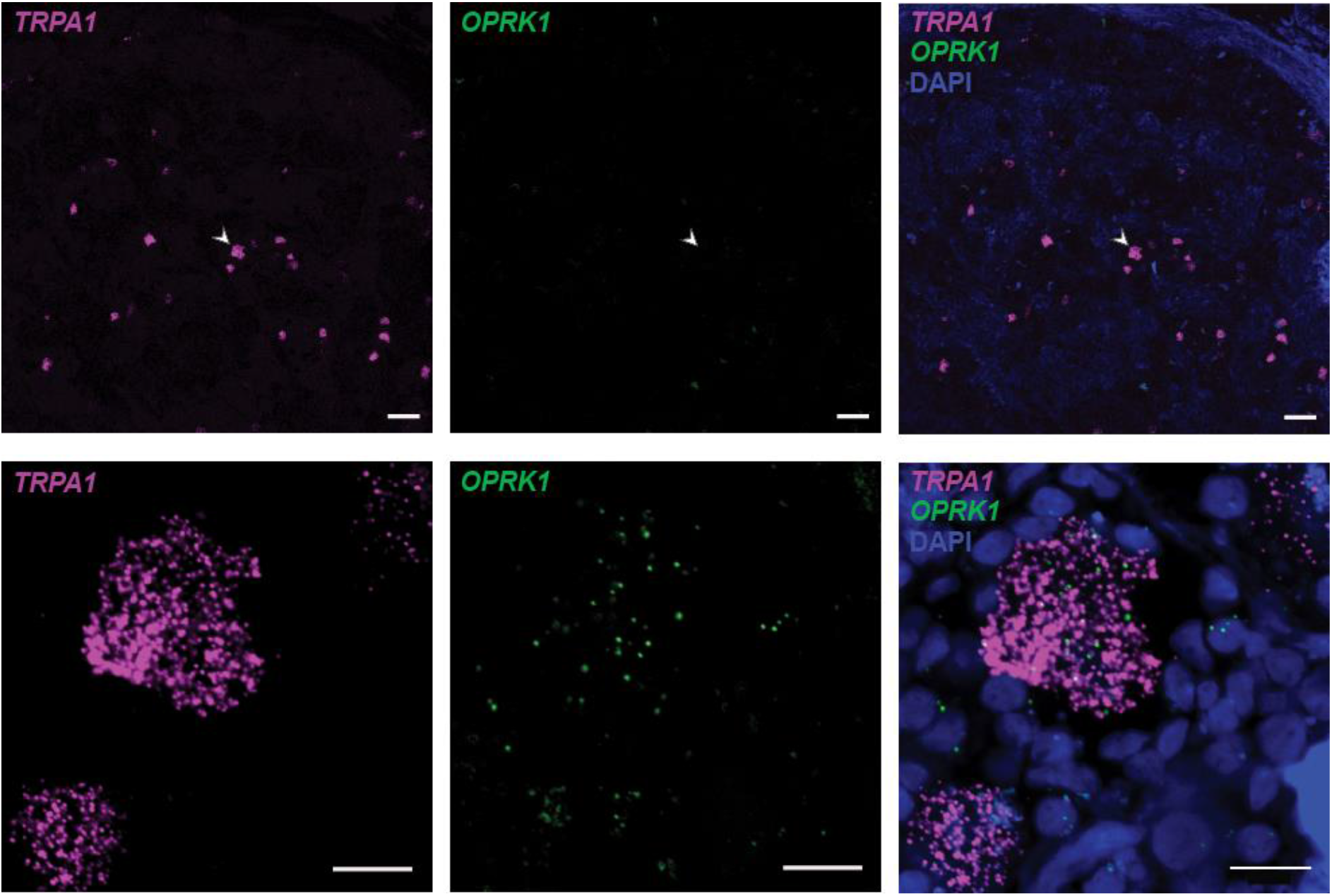
TRPA1 and KOR are expressed in human DRG neurons. *In situ* hybridization showing expression of *Trpa1* mRNA (purple) and *Oprk1* (green) in human DRG sections. Arrowhead marks a neuron co-expressing *Trpa1* and *Oprk1* at (top) low, 10x objective and (bottom) higher, 40x objective magnifications. Bars 100 μm (x10) and 20μm (x40).

### Constitutive KOR activity

Many class A GPCRs, such as opioid receptors show constitutive, agonist-independent receptor activity (Rosenbaum et al., 2009, Romero et al., 2012, Walwyn et al., 2016, Wang et al., 2007). To determine whether antinociception produced by TRPA1 inactivation is mediated by constitutive or agonist-evoked KOR activity *in vivo*, we sought inverse agonists and neutral antagonists that can distinguish between these two forms of inhibition. Inverse agonists inhibit both agonist evoked and spontaneous, constitutive receptor activity (Milligan, 2003) whereas neutral GPCR antagonists only inhibit agonist evoked activity.

We examined the mode of KOR antagonism produced by naloxone, 5’-GNTI, and naltrexone in KOR-expressing HEK293 cells using a cAMP luminescence assay. 5’-GNTI and naloxone dramatically enhanced the cAMP levels produced by forskolin (10 μM) stimulation of adenylate cyclase (Fig. 8a). This result demonstrated that KOR was constitutively active in HEK293 cells and that 5’-GNTI and naloxone act as inverse agonists. In contrast, naltrexone did not increase forskolin stimulated cAMP production and thus is not an inverse KOR agonist. As expected, the KOR agonist U50488 suppressed forskolin induced cAMP production completely (Fig. 8a).

**Figure 8.**
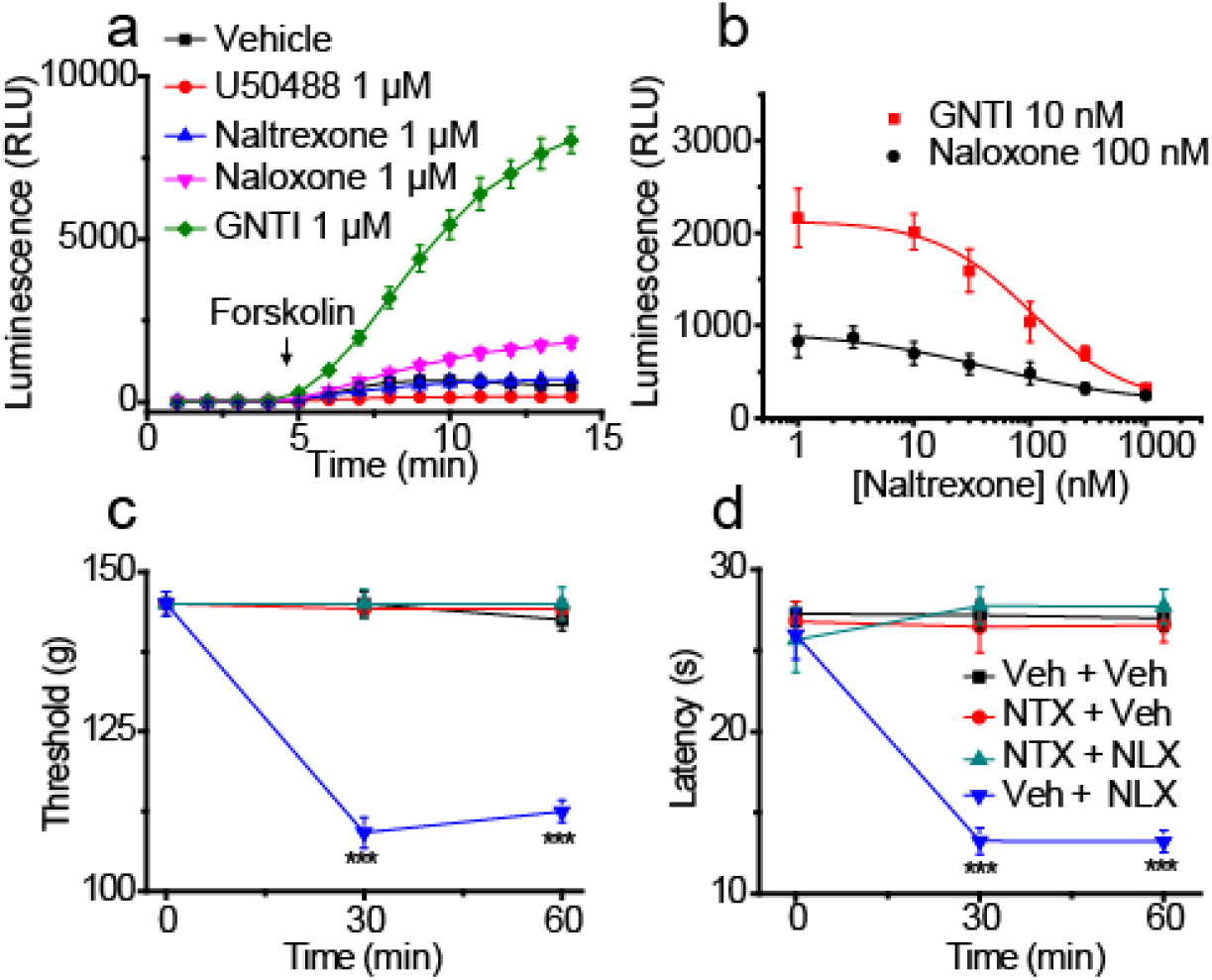
KOR is constitutively active *in vivo* and *in vitro*. (a) Effect of KOR ligands on forskolin (10μM) induced cAMP increases in HEK293 cells expressing a luminescent cAMP sensor and KOR. U50488 (KOR selective agonist) inhibited forskolin evoked cAMP production completely. Naloxone and 5’-GNTI acted as inverse agonists and enhanced cAMP production. Naltrexone is a neutral antagonist and had no effect. (b) Naltrexone concentration-dependently inhibited naloxone- and 5’-GNTI-augmented forskolin stimulated cAMP production. (c,d) Naloxone (2.5 mg/kg) normalized the behavioural sensitivity of Trpa1^-/-^ mice in the paw-pressure and cold-plate tests, whereas naltrexone (2.5 mg/kg) did not. ***P<0.001, compared to vehicle, 6 mice per group.

Since naltrexone behaved like a neutral antagonist (see also Wang et al., 2007), we examined whether it could inhibit the augmented cAMP response observed in the presence of the inverse agonists 5’-GNTI and naloxone. Naltrexone produced a concentration-dependent inhibition of 5’-GNTI- and naloxone-enhanced cAMP production, consistent with a neutral antagonist mode of action (Fig. 8b).

### Constitutive KOR activity in vivo

Having validated a KOR inverse agonist and a neutral antagonist we used these compounds to determine whether constitutive or agonist-dependent KOR activity was responsible for the sensory phenotype of *Trpa1^-/-^* mice. We compared the effects of naltrexone and naloxone on the behavioural sensitivity to noxious cold and mechanical stimulation (Fig. 8c, d). Strikingly, the neutral antagonist naltrexone was without effect on the behavioural sensitivity in the paw-pressure and cold plate tests, whereas the inverse agonist naloxone enhanced the sensitivity in both tests. These results suggest that constitutive KOR activity is responsible for the reduced sensitivities to noxious cold and mechanical stimulation in the absence of functional TRPA1. As neutral antagonists inhibit the pharmacological effects of both agonists and inverse agonists, we examined whether naltrexone could inhibit the effects of naloxone *in vivo*. The naloxone evoked reductions in paw withdrawal mechanical threshold and cold stimulation latency in *Trpa1^-/-^* mice were completely blocked by administration of naltrexone (Fig. 8c, d), as expected if naloxone was inhibiting constitutive KOR activity.

### Naloxone increases activity of AM- but not C-fibres in Trpa1^-/-^ mice

To gain insight into the effect of naloxone on sensory afferent nerve responses, we compared the activity of single units in ex-vivo skin-nerve preparations from *Trpa1^-/-^* and *Trpa1^+/+^* mice.

We challenged identified mechanosensitive single units with a single high force stimulus (15g, 150mN for 15s) either in the absence or presence of naloxone to avoid tachyphylaxis produced by repeated stimulation of fibres. The action potential frequency in both AM- and C-fibres was lower in *Trpa1^-/-^* than *Trpa1^+/+^* preparations (AM - Fig. 9a, d,e,m: CM – Fig. 9 g, j, k, n), in agreement with earlier findings (Kwan et al., 2009). In naloxone treated preparations, the frequency of mechanically evoked action potentials in AM-fibres from *Trpa1^-/-^* mice was raised to the level recorded in *Trpa1^+/+^* preparations (Fig. 9 c, f, m), whereas naloxone had no effect on AM-fibre responses in wild-type, *Trpa1^+/+^* preparations (Fig. 9 b, m). In contrast to its differential effect on AM-fibres, naloxone had no significant effect on the number of mechanically evoked C-fibre action potentials in either *Trpa1^-/-^* (Fig. 9 i, l, n) or *Trpa1^+/+^* preparations (Fig. 9 h,n), indicating that the TRPA1-KOR interaction primarily controls excitability in myelinated fibres.

**Figure 9.**
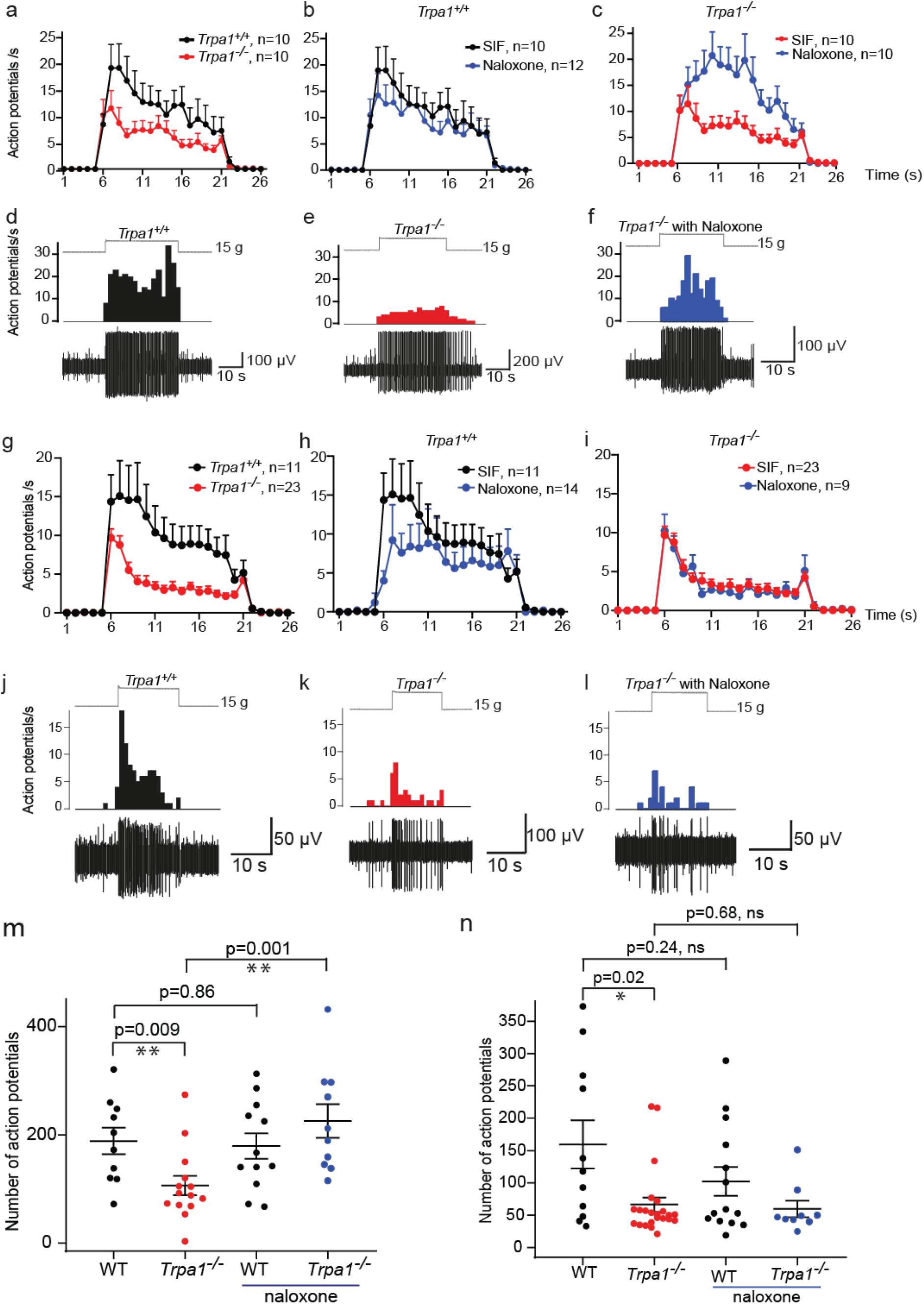
Naloxone increases the reduced activity of AM- but not C-fibres in *Trpa1^-/-^* mice. (a-f) responses of AM-fibres to 15 sec duration (5-20s), 15g mechanical stimuli. (a) Action potential frequency plots show lower firing rates in *Trpa1^-/-^* than *Trpa1^+/+^* fibres. (b) Naloxone has no effect on frequency in *Trpa1^+/+^* fibres but (c) increases the frequency in *Trpa1^-/-^* fibres. Examples of responses in fibres from (d) *Trpa1^+/+^* and (*e)Trpa1^-/-^* mice, and (f) *Trpa1^-/-^* fibre treated with naloxone. (g-l) C-fibre responses to same mechanical stimulation. (g) Lower rate of action potential firing in *Trpa1^-/-^* than *Trpa1^+/+^* fibres; examples shown for (j) *Trpa1^+/+^* and (k) *Trpa1^-/-^* fibres. Naloxone had no effect on firing rate in either *Trpa1^+/+^* or *Trpa1^-/-^* fibres. (m. n) Scatter plots of number of action potential evoked during the period of mechanical stimulation under different conditions in (m) AM-fibres and (n) C-fibres, illustrating the lower firing frequency in both classes of fibre and the increase in firing rate in AM-fibres in the presence of naloxone.

## DISCUSSION

TRPA1 antagonists have been investigated as potential analgesic drugs and are effective in a range of animal models of inflammatory and neuropathic pain where they reduce hypersensitivities to mechanical and thermal stimuli (for review see Koivisto et al., 2018). In conditions such as painful diabetic neuropathy, endogenous TRPA1 agonists are produced which are likely to contribute to ongoing pain by activating TRPA1. In this case, analgesia can at least in part be ascribed to direct inhibition of TRPA1 activation. However, it has been unclear how TRPA1 inhibition reduces mechanical and thermal hypersensitivities in other chronic pain conditions that are not driven by TRPA1 agonism (see Zygmunt and Hogestatt, 2014 for list of models tested). Our current study demonstrates that this results from a functional interaction between TRPA1 and KOR in sensory neurons.

The roles of TRPA1 in cold and mechanical sensation have been subject to controversy. Some studies failed to show cold-evoked responses in TRPA1 expressing cells including DRG neurons (Bautista et al., 2006, Kwan et al., 2006, Kwan and Corey, 2009, McKemy, 2005), while others reported that TRPA1 expression in cells conferred cold sensitivity (Chen et al., 2013, Jabba et al., 2014, Karashima et al., 2009, Moparthi et al., 2014, Sawada et al., 2007, Story et al., 2003, Memon et al., 2017). Our own studies have not found any direct link between TRPA1 expression and cold sensitivity in isolated DRG neurons. Furthermore, the majority of TRPA1-positive trigeminal ganglion and DRG neurons are not cold-sensitive (Jordt et al., 2004, Bautista et al., 2006, Karashima et al., 2009, Memon et al., 2017). There have also been conflicting reports for a role of TRPA1 in behavioural responses to cold. While we and others provided evidence that TRPA1 contributed to cold responses (Gentry et al., 2010, del Camino et al., 2010, Karashima et al., 2009), other groups reported that TRPA1 played no role (Knowlton et al., 2010, Bautista et al., 2006). These differences may, in part, be due to methodological differences, including the temperatures tested. Our data revealed that the sensitivity to noxious cold stimulation was reduced in both widely used lines of “global” *Trpa1^-/-^* mice, in mice with sensory neuron specific deletion of *Trpa1^-/^* and by pharmacological inhibition of TRPA1 in wild-type mice. However, the loss of cold sensitivity occurred over a restricted temperature range and we found no difference in behavioural responses between *Trpa1^-/-^* and wild-type mice at very cold temperatures that have been used in some other studies.

Although TRPA1 was proposed as a candidate mechanosensitive ion channel in inner-ear hair cells (Corey et al., 2004, Nagata et al., 2005) it was subsequently shown to have no essential mechanotransduction role (Kwan et al., 2006). There is also no evidence that TRPA1 is a mechanotransduction channel in either recombinant expression systems or in sensory neurons (see (Talavera et al., 2020). However, TRPA1 deletion or inhibition can affect mechanically evoked currents in isolated DRG neurons. TRPA1 deletion or pharmacological inhibition reduced slowly adapting currents in a subset of neurons (Vilceanu and Stucky, 2010) and the amplitude of intermediately adapting currents was reduced in *Trpa1^-/-^* DRG neurons (Brierley et al., 2011). Both effects would contribute to the reduced mechanically evoked sensory nerve firing observed in skin-nerve preparations from *Trpa1^-/-^* mice and after application of a TRPA1 antagonist to wild-type preparations (Kerstein et al., 2009, Kwan et al., 2009, Vilceanu and Stucky, 2010), as well as to the loss of mechanosensitivity in *Trpa1^-/-^* mice observed in this and earlier behavioural investigations (Andersson et al., 2009, Kwan et al., 2006, Zappia et al., 2017).

We have identified that the reduction in cold and mechanical sensitivities seen in *Trpa1^-/-^* mice was due to increased activity of opioid receptors in the periphery using a non-CNS penetrant analogue, naloxone methiodide, which was administered either systemically or by intraplantar injection where the effect was restricted to the injected paw. Furthermore, 5’-GNTI, which had a similar action to naloxone, is not CNS penetrant (Munro et al., 2012). A peripheral site of action of naloxone in *Trpa1^-/-^* mice differs from that operating in latent pain sensitization after tissue damage or inflammation where hypersensitivity is unmasked by inhibiting opioid activity in the spinal cord (Campillo et al., 2011, Corder et al., 2013, Walwyn et al., 2016).

Administration of opioid receptor subtype ligands identified KOR as the receptor sub-type responsible for the reduced sensitivities in *Trpa1^-/-^* mice. The findings that the levels of Oprk1 mRNA and expressed receptor protein as well as the endogenous KOR agonist dynorphin were unchanged in *Trpa1^-/-^* mice suggested that differences KOR activity could explain the reduced sensitivities. This differs from the mechanism proposed to underlie the opioid mediated antinociception in mice lacking Nav1.7. Here genetic ablation of Nav1.7 leads to increased levels of pro enkephalin (*Penk*) mRNA and protein (Minett et al., 2015), which signals via MOR and DOR, but not KOR (Pereira et al., 2018). Identification of naloxone and 5’GNTI as inverse agonists at KOR and naltrexone as a neutral antagonist allowed us to test the hypothesis that constitutive KOR activity was responsible for the loss of sensitivities in *Trpa1^-/-^* mice. Opioid receptors are known to be constitutively active (Romero et al., 2012, Walwyn et al., 2016, Wang et al., 2007) and constitutive receptor activity has been shown to regulate pain and pain remission associated with inflammation (Corder et al., 2013, Walwyn et al., 2016). Our *in vivo* results are consistent with a scheme in which TRPA1 regulates KOR activity. First, both inverse agonists, which inhibit constitutive KOR activity, reversed the loss of cold and mechanical in *Trpa1^-/-^* mice. Second, the neutral antagonist naltrexone had no effect alone but was able to inhibit the effects of naloxone. Despite intense investigations over decades, the mechanisms regulating constitutive activity of GPCRs remain unknown, but our results indicate that TRPA1 activity controls the constitutive activity of KOR. Administration of TRPA1 antagonists to wildtype mice recapitulated the phenotype of *Trpa1^-/-^* mice and the reduced cold and mechanical sensitivities were reversed by administration of naloxone. These acute effects of TRPA1 antagonists indicate that the regulation of KOR in sensory neurons is a dynamic process that does not result from either a compensatory effect during development or an effect on gene transcription/translation. The reduced behavioural sensitivity produced by acute TRPA1 inhibition also suggests that the channel displays some activity under normal physiological conditions.

A naloxone-reversible loss of cold and mechanical sensitivities was found with either global deletion of TRPA1 or after sensory neuron-specific deletion of TRPA1, demonstrating that the locus of action involved sensory neurons. This is consistent with our observed co-expression of *Trpa1* and *Oprk1* in DRG neurons. Studies using an *Oprk1-Cre* knock in allele to drive tdTomato showed that KOR was expressed in several sub-populations of mouse DRG neurons and found that about two-thirds of *Oprk1-Cre* labelled sensory neurons are nociceptive with 50% expressing TRPV1 (Snyder et al., 2018). Our ISH results showed a similar degree of co-expression with *Trpa1* expressed in about 40% of *Oprk1*-positive DRG neurons. The percentage of KOR expressing mouse DRG neurons is only about 10-11%, similar to the percentage seen in rats (Ji et al., 1995, Schafer et al., 1994, Zhang et al., 1998); however, altered activity in these neurons can have a marked analgesic effect. For example, the KOR-specific agonist U50,488 has a local analgesic action when injected into the paw (Auh and Ro, 2012, Moon et al., 2016) and peripherally restricted KOR agonists are analgesic (Beck and Dix, 2019, Kivell and Prisinzano, 2010, Vanderah, 2010).

A previous study demonstrated that mechanosensitive C-fibres in *Trpa1^-/-^* mice displayed a reduced impulse frequency in response to mechanical stimulation at all forces, while A-mechanosensitive (AM) fibres showed reduced action potential firing but only with higher force stimuli (Kwan et al., 2009). We found that naloxone increased the impulse rate in AM-fibres but not C-fibres in *Trpa1^-/-^* mice. This is consistent with our ISH/immunostaining results and those of Snyder et al. (Snyder et al., 2018) which showed that *Oprk1* was expressed together with CGRP and a neurofilament marker in small to medium diameter neurons.

Our results also demonstrate that increased opioid receptor activity can explain the reduced sensitivities of *Trpa1^-/-^* mice in chronic pain models. Here the development of hypersensitivities following nerve ligation or inflammation occurred on elevated baseline paw pressure thresholds and cold plate latencies. Increases in cold and mechanical sensitivities were noted in both models, but the sensitivities remained lower in *Trpa1^-/-^* than in *Trpa1^+/+^* mice. Strikingly, administration of naloxone increased cold and mechanical sensitivities in *Trpa1^-/-^* mice to the same levels seen in hypersensitive *Trpa1^+/+^* mice indicating that opioid receptor activity masked the full appearance of hypersensitivities.

TRPA1 antagonists reduce established sensory hypersensitivities in experimental models of inflammation and neuropathy in wild type mice (see reviews by (Koivisto et al., 2018, Moran and Szallasi, 2018)). The finding that naloxone inhibited the anti-hyperalgesic effects of the TRPA1 antagonist A967079 in both inflammatory and neuropathic conditions demonstrates that in these mouse models of chronic pain the effects of TRPA1 antagonism can be explained by altered activity of opioid receptors.

GPCR regulation of TRP channel activity is well established (see Veldhuis et al., 2015), but the concept that the activity of TRP channels can regulate GPCR signalling is novel. We speculate that this is likely to operate in other types of neurons as well as non-neuronal cells throughout the body. Apart from our study there are few other reports of TRP channels modulating GPCR function. Two studies have demonstrated that TRPV1 activation can inhibit agonist evoked internalization and desensitization of MOR (Basso et al., 2019, Scherer et al., 2017), and activation of TRPV4 channels has been demonstrated to inhibit internalization of angiotensin II, type 1 receptors (Zaccor et al., 2020). Although this type of regulation may occur for TRPA1 and KOR, such a mechanism cannot explain our finding that constitutive KOR activity is increased in the absence of TRPA1 activity.

The mechanism by which the presence or absence of functional TRPA1 channels regulates constitutive activity of KOR remains to be elucidated. Surprisingly, little is known about the cellular and molecular mechanisms controlling the constitutive activity of GPCRs. Our results are consistent with a scheme where a normal level of TRPA1 activity reduces the constitutive activity of KOR (Fig 10a) and this repression is lost when TRPA1 is silenced either genetically or pharmacologically (Fig. 10b). TRPA1 regulation of KOR activity could be due to an activity-dependent structural interaction between the proteins or result from some downstream functional effects of TRPA1 activity, and this is an area for future investigations.

**Figure 10.**
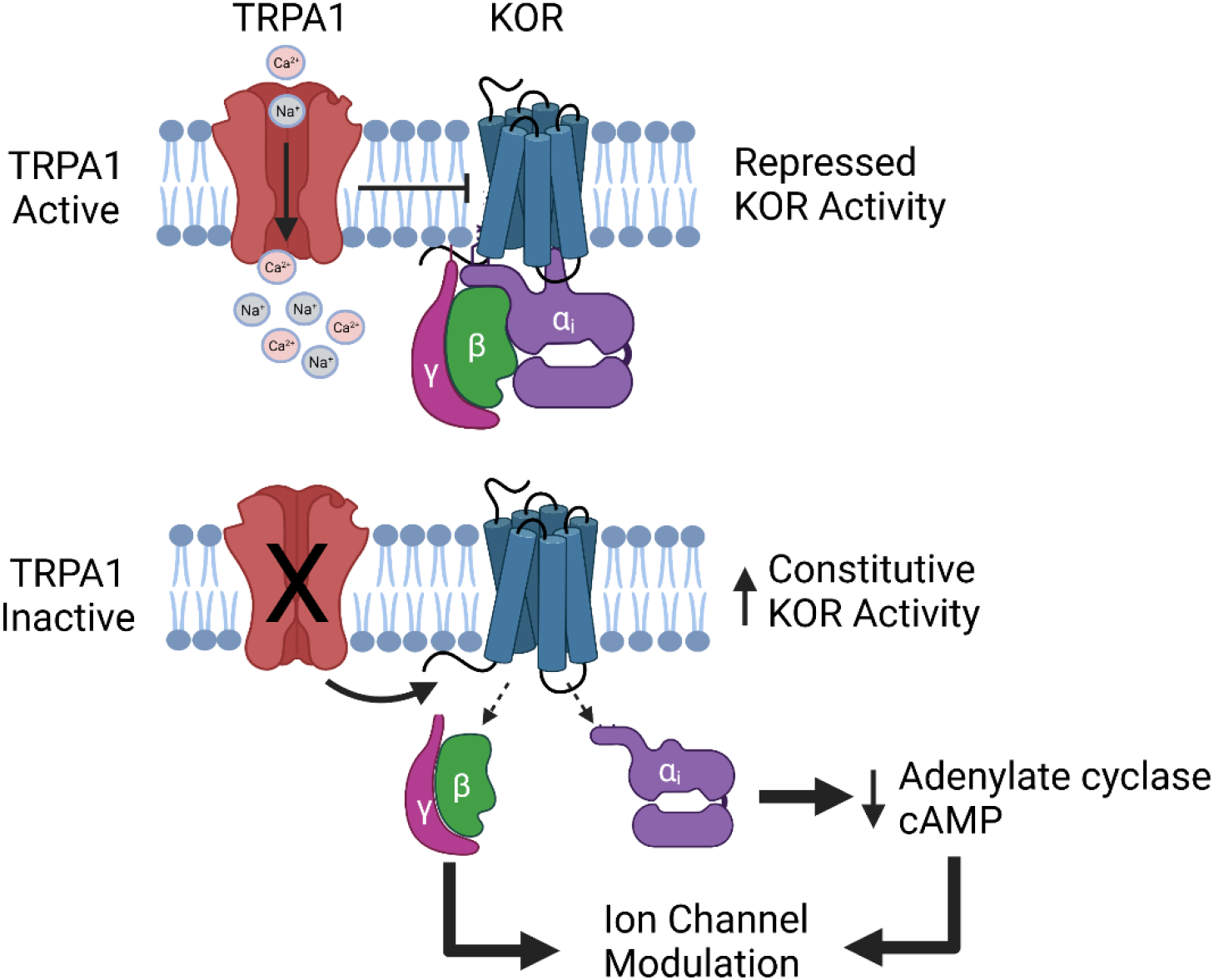
Schematic diagram of mechanism for TRPA1 regulation of KOR. (a) Normal TRPA1 activity represses KOR by a structural or functional interaction. Genetic or pharmacological silencing of TRPA1 relieves the inhibition leading to increased constitutive KOR activity which reduces sensory neuronal excitability by modulating the activities of e.g. Kir, HCN, K2P channels. Created with BioRender.com.

How activation of KOR in peripheral sensory neurons reduces action potential firing is also unclear. One possible mechanism is by Gβγ potentiation of GIRK1-4 (Kir3.1-3.4) potassium channel activity. Although this mechanism occurs post-synaptically in the spinal cord for MOR and DOR (Marker et al., 2005) there is little to no current evidence that this operates for KOR in sensory neurons. Several studies failed to detect GIRK-mediated currents or responses in mouse DRG neurons (Kanjhan et al., 2005, Nockemann et al., 2013, Saloman et al., 2016). However, expression of GIRK1 and GIRK 2 have been reported in mouse DRG neurons (Saloman et al., 2016, Zeisel et al., 2018) and KOR modulation of GIRK channel activity remains a possibility. cAMP modulation of HCN channels is another candidate mechanism (Ingram and Williams, 1994). KOR activation will lead to inhibition of adenylate cyclase and reduced cAMP production in DRG neurons (Berg et al., 2011, Jamshidi et al., 2015). This can result in decreased activation of HCN channels, which will reduce neuronal excitability and repetitive firing in DRG neurons (Emery et al., 2012). Altered cAMP/PKA-mediated regulation of other voltage gated sodium and potassium channels could also modulate neuronal activity (England et al., 1996, Evans et al., 1999, Gold et al., 1996, Gold et al., 1998). In addition, there is evidence that cAMP regulation of two pore potassium channels (TREK-1) in sensory neurons contributes to MOR-mediated analgesia (Devilliers et al., 2013) and this may also operate for KOR.

The need for novel, effective analgesic drugs with minimal or no side effects or abuse liability has been heightened by the opioid crisis (Schuchat et al., 2017, Coussens et al., 2019). Our finding that inhibition of TRPA1 can result in constitutive KOR activity and consequent analgesia suggests that this mechanism could be exploited for effective treatments in humans as transcripts for TRPA1 and KOR are co-expressed in human DRG neurons. A TRPA1 antagonist will act directly to inhibit neuronal activity or hypersensitivity driven by endogenous molecules that activate TRPA1, while TRPA1 inactivation would also act indirectly to promote KOR activity and thereby produce a more general analgesic profile. Our finding that the constitutive activity of GPCRs is influenced by the activity of a TRP channel has potentially wider implications as TRP channels are co-expressed with GPCRs in cells throughout the body, where ion channel activity could influence GPCR signalling.

## MATERIALS AND METHODS

### Behavioural studies

All animal studies were performed according to the UK Home Office Animal Procedures Act (1986) after ethical review and approval by the King’s College London Animal Welfare and Ethical Review Body. Data shown are from male and female *Trpa1^-/-^, Trpa1^+/-^* and *Trpa1^+/+^* littermates kindly provided by Kelvin Kwan and David Corey (Harvard Medical School) (Kwan et al., 2006) and David Julius (University of San Francisco) (Bautista et al., 2006). Both strains were back crossed onto the C57BL/6J strain (Harlan/Envigo, Blackthorn, UK) for >12 generations. Unless stated otherwise the data presented are for the Kwan/Corey *Trpa1^-/--^* mice. There were no sex differences identified in any behavioural test and so data from male and female mice have been amalgamated. Thermal sensitivity was assessed using a commercially available hot- and cold-plate (Ugo Basile, Milan). Paw withdrawal latencies were determined with the plate set at a chosen temperature (in the range 2-53 °C). Mice were lightly restrained and each hind paw in turn was placed onto the surface of the plate. The end point was taken as the withdrawal of the paw and recorded as the withdrawal latency for the ipsilateral and the contralateral paw with a cut-off latency of 30s. Mechanical sensitivity was assessed in lightly restrained mice by measuring paw withdrawal thresholds to an increasing mechanical force applied to the dorsal surface of the paw using an Analgesymeter (Ugo-Basile, Milan, Italy) with a cut-off force of 150g.

Neuropathic pain was induced by partial nerve ligation (Seltzer et al., 1990). The sciatic nerve in the left leg was exposed at high thigh level through a small incision, cleared of surrounding connective tissue and 1/3 to 1/2 of the dorsal nerve tightly ligated with a 7-0 silk suture. Inflammatory hypersensitivity was induced by intraplantar injection of 10-15μl of Complete Freund’s Adjuvant (CFA).

For the abdominal constriction test, an intraperitoneal injection of dilute phenyl-*p*-benzoquinone (0.02 %, 0.2 ml, i.p) was used to induce a nociceptive stereotyped behaviour. The intensity of pain was quantified by counting the number of abdominal stretches that occurred during a 20 min observation period. The time spent on nocifensive behaviour (paw lifting, biting, licking, shaking) following an intraplantar injection of AITC (20 μl of 75 mM) was quantified over a 10 min period. HC-030031 was from TosLab (Ekaterinburg, Russia), A967079 was from Manchester Organics (Runcorn, UK). All other drugs were from Sigma-Aldrich (Gillingham, UK) or Tocris (Bristol, UK). Treatments were randomly allocated to mice in each cage and the operator was blinded for the outcome assessment and experiments conducted according to the ARRIVE guidelines.

### [cAMP]_i_-measurements

Human derived HEK293 cells (Invitrogen, Inchinnan, UK, R75007) stably transfected with cAMP GloSensor™ (pGloSensor-22F; Promega, Southampton, UK) and mouse *Oprk1* (Origene, Rockville, USA) were grown in DMEM (Gibco) AQ supplemented with penicillin (100 U/ml), streptomycin (100 μg/ml), FBS (10%), G418 (0.5mg/ml) and hygromycin (200 μg/ml) and were studied in 96-well plate assays. Cells were loaded with GloSensor™ cAMP reagent (2mM) for 1-2 hours at 37°C and luminescence monitored using a Flexstation 3 (Molecular Devices, San Jose, USA) at 25°C. Cells were incubated with opioid receptor ligands for 10 minutes before addition of forskolin (10 μM) to stimulate cAMP production. Luminescence was monitored for 15 min following addition of forskolin. Data are shown as mean ± SEM of triplicate wells.

### RNA extraction

RNA was extracted from DRGs of C57Bl6/J wild type (Harlan, UK) and *Trpa1^-/-^* mice (Kwan et al., 2006), age 20 weeks, and stored until use at −20°C in RNA later (Thermo Fisher Scientific, Waltham, USA). The tissue was homogenized in Trizol (Invitrogen) and samples incubated in chloroform (Sigma-Aldrich, Gillingham, UK) in 5Prime Phase Lock Gel tubes (Quantabio, Beverley, USA). Tubes were spun for 15 minutes and the RNA in the aqueous phase was precipitated with 100% ethanol and transferred into an RNeasy column (Qiagen). 30μl of RNA per sample were collected and the concentration and purity were determined using a Nanodrop 1000 (Thermo Fisher Scientific, Waltham, USA).

### Quantitative RT-PCR

Total RNA from the DRGs was reverse transcribed using a High-Capacity RNA-to-cDNA Reverse Transcription Kit (Applied Biosystems, Waltham, USA) according to the manufacturer’s instructions. The cDNA from wild type and *Trpa1^-/-^* DRGs was targeted using custom TaqMan gene expression assays (Applied Biosystems) which included a mix of forward and reverse primers, as well as a FAM-labelled TaqMan MGB probe. Four different TaqMan gene expression assays were used; *Gapdh* (Mm99999915_g1), *Oprk1* (Mm01230885_m1) and *Trpa1* (Mm00625268_m1). The cDNA and primer sequences of all genes are stated in Table 1. *Trpa1* cDNA was targeted at two different regions, the channel’s S5 and S6 transmembrane domains (exons 21-25), which are deleted in the *Trpa1^-/-^* mice genome (Kwan et al., 2006) and the Ankyrin repeat region (exons 7-8), which is present in both wild type and *Trpa1^-/-^* animals genome.

**Table 1.**
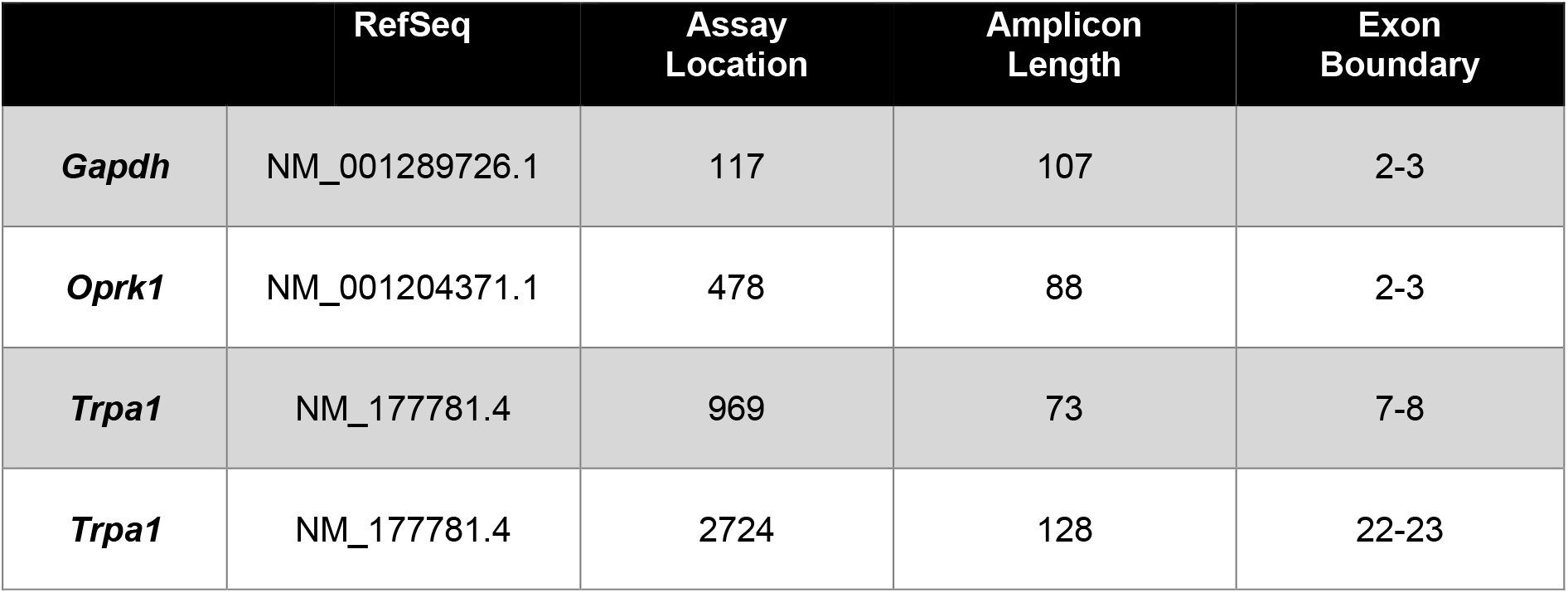
TaqMan gene expression assays (Applied Biosystems) used for qRT-PCR.

|  | RefSeq | Assay Location | Amplicon Length | Exon Boundary |
| --- | --- | --- | --- | --- |
| **Gapdh** | NM_001289726.1 | 117 | 107 | 2-3 |
| **Oprk1** | NM_001204371.1 | 478 | 88 | 2-3 |
| **Trpa1** | NM_177781.4 | 969 | 73 | 7-8 |
| **Trpa1** | NM_177781.4 | 2724 | 128 | 22-23 |

qRT-PCR amplification was performed with a LightCycler 480 (Roche Life Science, Welwyn Garden City, UK) using 45 cycles (initial heat denaturation 95°C/10min; per cycle 95°C/10s denaturation, 60°C/30s annealing, 72°C/1s extension; final cooling 40°C/30s, fluorescence measured at 521nm). cDNA samples were amplified in quadruplets (four technical replicates) per candidate gene and 10 biological replicates (samples).

Data were analysed using LightCycler 480 Software, version 1.5.1 (Roche Life Science). The mean threshold cycle (CT) for each quadruplet per sample was used for further analysis. qRT-PCR data were presented as expression fold change following the comparative 2^-ΔΔCT^ method (Schmittgen and Livak, 2008) with each gene (*Oprk1* and *Trpa1*) normalized to the housekeeping gene, *Gapdh*.

### RNAscope in situ hybridization

#### Mouse DRGs

A standard manual multiplex fluorescent assay was conducted (Wang et al., 2012). DRGs were dissected from wild type and *Trpa1^-/-^* mice and fixed in 4% paraformaldehyde (Thermo Fisher Scientific). Samples were washed with phosphate buffered saline, dehydrated in ethanol followed by xylene, and embedded in paraffin. 6μm sections were cut, air dried and baked for 1 hour at 60°C. The paraffin was removed with a series of xylene and 100% ethanol washes. The sections were heated for 5min at 60°C and then covered with Hydrogen Peroxide for 10min at room temperature. Slides were placed in 100°C Target Retrieval solution (Advanced Cell Diagnostics, Newark, USA) for 15min, and then covered with RNAscope Protease Plus (Advanced Cell Diagnostics) and incubated for 30min at 40°C.

Samples were processed using the RNAscope multiplex fluorescent kit v2 (Advanced Cell Diagnostics). RNAscope probes (Table 2) were diluted in probe diluent (1:50 dilution) and stored at 4°C before use. Sections were covered with a mixture of the two probes (*Trpa1* and *Oprk1*) and hybridized in a HybEZ oven for 2 hours at 40°C. After the hybridization step, AMP1, AMP2, AMP3 signal amplification molecules were applied to the sections for 30min each.

**Table 2.** RNAscope probes (Advanced Cell Diagnostics) used for *in situ* hybridization of mouse DRGs.

|  | RefSeq | Targeting Region | Exon Boundary |
| --- | --- | --- | --- |
| **Mm-Oprk1-C3** | NM_001204371.1 | 256-1457 | 2-3 |
| **Mm-Trpa1-C2** | NM_177781.4 | 77-1047 | 1-7 |

**Table 3.** RN<u>Ascope</u> probes (Advanced Cell Diagnostics) used for *in situ* hybridization of human DRGs.

|  | RefSeq | Targeting Region | Exon Boundary |
| --- | --- | --- | --- |
| Hs-Oprk1-C2 | NM_000912 | 2172-3208 | 4 |
| Hs-Trpa1-O1 | NM_007332.2 | 3148-4496 | 25-27 |

The probes were assigned different channels (*Oprk1*-C3 and *Trpa1-C2*) and the HRP signal for these channels was developed separately for 15min at 40°C using the RNAscope HRP-C1 and HRP-C2 reagents (Applied Cell Diagnostics). Following HRP development, TSA Plus Cyanine3 (Cy3, for *Trpa1*) and Cyanine 5 (Cy5, for *Oprk1*) fluorophores (Perkin Elmer, 1:1500 dilution) were applied for 30min at 40°C. The HRP for each channel was blocked using the RNAscope HRP blocker. The sections were counterstained with DAPI for 30sec at room temperature, mounted with Vectashield mounting media (VWR) and stored at 4°C.

##### Human DRGs

Fresh frozen human DRGs from 3 individuals without chronic pain, aged over 60 years old, were provided by the Netherlands Brain Bank (NBB) to Eli Lilly & Co Research Centre, UK. The tissue is distributed to researchers with an anonymized detailed medical record of the donor. Human DRGs sourced by NBB were dissected during autopsies and immediately frozen. Upon arrival in the UK, the tissue was embedded in optimal cutting tissue component (OCT) (Sakura) and stored at −80 ° C. The DRGs were sectioned at 10μm using a Cryostat (Leica Biosystems), at −20 ° C. Approximately 2-3 sections were mounted on each Superfrost Plus slide (Thermo Fisher Scientific). All sections were air-dried at −20 ° C for 1 hour and afterwards stored at −80 ° C.

Before the initiation of the RNAscope assay, the sections were removed from −80 °C and fixed in 4% paraformaldehyde (PFA) at 4 ° C. PFA was washed and the sections were dehydrated by a series of increasing concentration ethanol washes. The sections were then incubated in Hydrogen Peroxide, followed by a Protease IV (Advanced Cell Diagnostics) incubation step for 30 minutes at room temperature. The samples were then processed using the protocol described above for mouse DRGs.

Images from all in situ hybridization experiments were captured using an LSM710 confocal microscope (Zeiss) and ZEN2010 software (Zeiss).

#### Combined Immunostaining and in situ hybridization

After completion of the in situ hybridization assay, sections were washed with TBST wash buffer (10% Tween20 in 1xTBS buffer) and incubated in 10% sheep serum (Sigma Aldrich) in TBS-1%BSA (Sigma Aldrich), for 30min at room temperature. Primary antibodies directed against NF200 (monoclonal anti-mouse IgG, 1:400 dilution, Sigma Aldrich, SAB4200747) and CGRP (polyclonal anti-rabbit IgG, 1:1,000 dilution, Enzo Life Sciences, BML-CA1134) were applied for 2 hours at room temperature. Following a wash, secondary antibodies (Alexa 594 goat anti-mouse IgG (H+L), 115-585-003, RRID: AB_2338871; Alexa 488 donkey anti-rabbit IgG, 1:1,000 dilution, 711-545-152, BRID AB_2313584, Jackson ImmunoResearch, Ely, UK) were applied to the sections for 30min at room temperature. The sections were washed, counterstained with DAPI (Advanced Cell Diagnostics), mounted using Vectashield (VWR) and stored at 4°C in the dark.

#### Microscopy and data analysis

Images from the *in situ* hybridization and combined in situ hybridization/immunofluorescence experiments were obtained using an LSM710 confocal microscope (Zeiss) and ZEN2010 software (Zeiss). To detect all markers, three different laser lines (405nm, 514nm, 633nm) were used with constant laser intensity and other scanning parameters for all sections. Single focal plane images of all channels were taken for in situ hybridization experiments using 10x, 20x and 40x objectives. Z-stacks of lμm/step were used to count the number of *Oprk1* and *Trpa1* positive cells in mouse DRGs. 50 z-stacks were analysed for each group (3 wild type and 3 *Trpa1^-/-^* mice). For combined *in situ* hybridization/ immunofluorescence experiments, 20x single focal plane pictures were taken. Similarly, only 10x single focal plane pictures were obtained for the *OPRK1* and *TRPA1 in situ* hybridization study in human DRGs,.

All pictures were processed in Fiji (ImageJ) and NIS-Elements (Nikon). Two different types of analysis were performed. Fiji (ImageJ) was used to count the number of positive cells for each transcript (*Oprk1* and *Trpa1*) and the total number of cells per z-stack. The criteria for the selection of the positive cells were the puncta that depict the mRNA molecules, and the visible cell outline. The cell diameter analysis for *Oprk1* and/or *Trpa1* positive cells was performed using the NIS-Elements software (Nikon). Data show the mean percentages of positive cells per animal for each transcript ±SEM, for each cell area.

#### Protein expression analysis

Protein was extracted from mouse (*Trpa1^+/+^* and *Trpa1^-/-^*) DRGs by phenol-chloroform extraction. DRG protein was obtained from samples used for RNA extraction. DNA was isolated from the lower phase of all samples and the supernatant was used for protein isolation. Proteins were precipitated, washed and then solubilized in 1% SDS (Sigma-Aldrich) and stored at −20°C before analysis. The protein concentrations were estimated using the Braford assay using bovine serum albumin standards (Thermo Fisher Scientific) with absorbance measured at 595nm.

#### Western blots

30-50μg of proteins from mouse DRGs were boiled for 5min in sample buffer at 95°C, and loaded on a 10% concentrated 1mm gel (Protogel; Sigma-Aldrich) and subjected to electrophoresis (100V for 90min). Proteins were transferred to a PVDF membrane (60V for 90min) and the membrane placed into blocking solution (5% milk powder in 0.05% PBS-T) for 30min at room temperature. Recombinant rabbit monoclonal antibody for OPRK1 (ab183825, Abcam) was added to the membrane at a 1:1000 dilution and incubated overnight at 4°C. The membrane was washed (3×5min in 0.05% PBS-T) and then blocked in 5% milk in 0.05% PBS-T for 30min. Blocking solution was removed, and the membrane incubated with an Alexa Fluor 680 goat anti-rabbit IgG secondary antibody (Invitrogen, 1:5000 dilution) in the dark for 1 hour at room temperature. The membrane was then washed (3×5min with 0.05% PBS-T) and visualized using an Odyssey machine (LI-COR). Final image adjustments were done using the Image Studio Lite software (LI-COR). The same protocol was used for β-actin detection using a rabbit polyclonal primary antibody (ab8227, Abcam, 1:1000 dilution). As noted by the manufacturer, the recombinant rabbit monoclonal antibody which recognizes a 300aa sequence at the C-terminus of the OPRK1, detects a glycosylated OPRK1 band of approximately 58kDa, whereas the predicted molecular weight for OPRK1 is 43kDa.

##### ELISA Assays

Protein levels were measured using ELISA kits for dynorphin (E03D0267 Blue Gene, Shanghai) and KOR (SED043Mu, Cloud Clone Corp, Houston) according to the manufacturers’ instructions.

##### Skin-nerve electrophysiological recordings

The isolated saphenous skin-nerve preparation and single fibre recording technique was used to record from the terminals of primary afferent neurons as described previously (Koltzenburg et al., 1997, Reeh, 1986). Recordings were carried out from 8-12 week old adult *Trpa1^-/-^* and *Trpa1^+/+^* mice. Briefly the saphenous nerve and hind-paw hairy skin were dissected free and immediately placed in warm (32°C) oxygenated synthetic interstitial fluid (SIF) containing (in mM); 108 NaCl, 3.5 KCl, 0.7 MgSO_4_, 26.2 NaHCO_3_, 1.65 NaH_2_PO_4_, 9.6 Na-gluconate, 5.55 glucose, 1.53 CaCl_2_, buffered to pH 7.4 by bubbling carbogen (95% O_2_ and 5% CO_2_). The skin was pinned down, corium side up and the saphenous nerve was placed on a mirrored platform covered in paraffin oil in a separate chamber containing a SIF-paraffin oil interface. The nerve was desheathed and teased into thin filaments, which were placed onto a gold electrode. Electrical signals were recorded using the DAM80 differential amplifier (World Precision Instruments), band-pass filtered at 300 Hz and 10 kHz and digitized and stored on a computer using the Micro1401 data acquisition unit and SPIKE2, version 8 software (Cambridge Electronic Design). Action potentials were also visualized using an oscilloscope (Gould, 420 series).

Receptive fields of mechanosensitive primary afferents were identified using a blunt glass rod and thereafter stimulated with brief electrical pulses (1ms) (Digitimer, DS2) to determine the fibre’s conduction velocity. For the purpose of this study only mechanically sensitive unmyelinated C fibres (CM) which conducted below 1.2 m/s and high threshold thinly myelinated Aδ fibres (AM) conducting between 1.2 and 10m/s were investigated, as these population of neurons are known to express TRPA1(Barabas et al., 2012, Kwan et al., 2009). Mechanical sensitivity of single A and C fibre nociceptors were assessed by applying a suprathreshold 15s duration sustained mechanical force of 15g delivered by a computer driven, feedback controlled stimulator (De Col et al., 2012), which controlled a plastic stylus with a tip diameter of 1mm. Fibres were classified as AM and CM if they exhibited slowly adapting responses to the 15g sustained force.

In a separate set of experiments, the naloxone hydrochloride (Sigma) was added to the circulating SIF (final concentration of 10μM from stock solution of 10mM made up in water) to determine its effect on mechanical sensitivity of A and C fibre nociceptors. Mechanically evoked action potentials were analysed offline using the SPIKE2 software (Cambridge Electronic Design). Action potentials were discriminated based on their amplitude and waveform and subsequently quantified.

##### Statistics

All data are expressed as mean ± SEM. The results of behavioural experiments were analyzed by repeated measures ANOVA followed by Tukey’s, Dunnett’s or Sidak’s *post hoc* tests or unpaired t-test, as appropriate. ELISA results were analyzed by unpaired t-tests. For the skin-nerve studies, differences in mechanically evoked action potentials were analysed using Levene’s test of equality followed by a Mann-Whitney *U*-test. Differences were considered significant at p<0.05.

## ACKNOWLEGEMENTS

We thank Drs. Kelvin Kwan and David Corey, and David Julius for the gifts of their respective strains of *Trpa1^-/-^* mice, Dr. Adrian Mogg for supervising ES at Eli Lilly & Co., Dr. Lisa Broad for support (ES) and providing access to human DRGs and Dr. Maria Levitin for human DRG sections.

## AUTHOR CONTRIBUTIONS

Conceptualization, S.B., D.A.A. C.G. and E.S.

Methodology, C.G. E.S. and C.S.

Formal Analysis, S.B., D.A.A. C.G, E.S, and N.V.

Investigation, C.G., E.S. and N.V.

Resources, C.S.

Writing – Original Draft, S.B., D.A.A, C.G., E.S. and N.V.

Writing – Review and Editing, C.S.

Funding Acquisition, S.B., D.A.A. and C.S.

Supervision, D.A.A. and S.B.

## DATA AND MATERIALS AVAILABILITY

All data are available in the main text or the supplementary materials

---

**Supplementary Figure 1.**
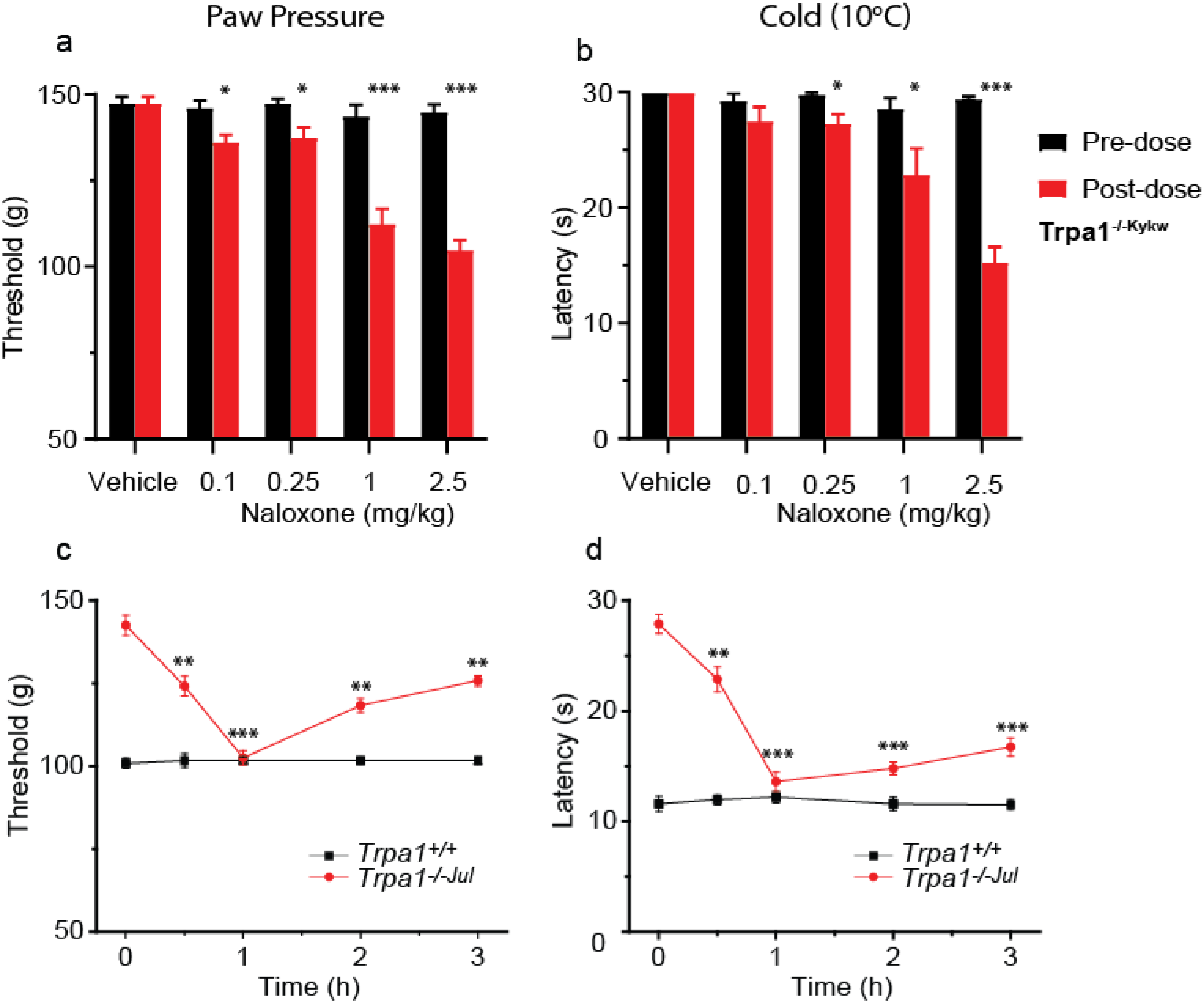
Naloxone dose-dependently increases sensitivities in *Trpa1^-/-Kykw^* mice and normalizes mechanical and cold sensitivities in *Trpa1^-/-Jul^* mice. Increase in mechanical (a) and cold (b) sensitivities in *Trpa1^-/kykw^* mice 1h after administration of 0.1, 0.25, 1 or 2.5mg/kg naloxone. Naloxone (2.5mg/kg, i.p.) reduces paw-pressure withdrawal threshold (c) and cold-plate latency (d, 10°C) in *Trpa1*^-/-Jul^ but not *Trpa1^+/+^* mice. *P<0.05, **P < 0.01, ***P <0.001. Data are mean ± SEM, 6 mice per group.

